# Actin filaments couple the protrusive tips to the nucleus through the I-BAR domain protein IRSp53 for migration of elongated cells on 1D fibers

**DOI:** 10.1101/2022.05.20.492840

**Authors:** Apratim Mukherjee, Jonathan E. Ron, Hooi Ting Hu, Tamako Nishimura, Kyoko Hanawa-Suetsugu, Bahareh Behkam, Nir S. Gov, Shiro Suetsugu, Amrinder S. Nain

**Affiliations:** Department of Mechanical Engineering, Virginia Tech, Blacksburg, 24061, USA; Department of Chemical and Biological Physics, Weizmann Institute of Science, Rehovot, 7610001, Israel; Data Science Center, Nara Institute of Science and Technology, Ikoma 630-0192, Japan; Center for Digital Green-innovation, Nara Institute of Science and Technology, Ikoma 630-0192, Japan; Graduate School of Brain Science, Doshisha University, Kyotanabe, Kyoto 610-0394, Japan

**Keywords:** IRSp53, cell forces, protrusions, ECM nanofibers, membrane curvature, stick-slip migration, actin, retrograde flow

## Abstract

The cell migration cycle proceeds with shaping the membrane to form new protrusive structures and redistribution of contractile machinery. The molecular mechanisms of cell migration are well-studied in 2D, but membrane shape-driven molecular migratory landscape in 3D fibrous matrices remains poorly described. 1D fibers recapitulate 3D migration, and here, we examined the role of membrane curvature regulator IRSp53 as a coupler between actin filaments and plasma membrane during cell migration on suspended 1D fibers. Cells attached, elongated, and migrated on the 1D fibers with the coiling of their leading-edge protrusions. IRSp53 depletion reduced cell-length spanning actin stress fibers, reduced protrusive activity, and contractility, leading to uncoupling of the nucleus from cellular movements. Using a theoretical model, the observed transition of IRSp53 depleted cells from rapid stick-slip migration to smooth, and slower migration was predicted to arise from reduced actin polymerization at the cell edges, which was verified by direct measurements of retrograde actin flow using speckle microscopy. Overall, we trace the effects of IRSp53 deep inside the cell from its actin-related activity at the cellular tips, thus demonstrating a unique role of IRSp53 in controlling cell migration in 3D.

## 1. Introduction

Cell migration has been extensively studied using anisotropic featured 2D and 3D matrix systems. Flat 2D migration assays have limited physiological relevance, while 3D gels are intrinsically heterogeneous and complex ^[1]^, which makes it difficult to understand the role of each constituent fiber in cell migration. Interestingly, cell migration was shown to be recapitulated through the use of narrow 1D microprinted lines on a 2D surface ^[2–5]^, which are essentially flat and are not fibers. We have previously shown that suspended, 1D synthetic fibers of sub-micron diameters coated with extracellular matrices (ECM) proteins result in protrusive, contractile, and migratory behavior that are sensitive to fiber curvature ^[6–11]^.

Cell attachment and migration on the suspended 1D fibers begins with characteristic protrusions that *coil* around the fiber axis ^[8,12]^. In 2D and 3D, filopodial protrusions are mediated by the actin cytoskeleton and the inverse Bin/Amphiphysin/Rvs (I-BAR) domain-containing proteins ^[13–15]^. The I-BAR domain proteins are crucial regulators of cell membrane curvature for protrusion formation, membrane trafficking, and cell migration, including cancer metastasis ^[16–20]^. Structurally, the intrinsically convex shaped surface of the I-BAR proteins have been demonstrated to drive the extension of cell protrusions such as thin, “finger-like” filopodia and broader, “sheet-like” lamellipodia ^[21–23]^. In support of these experimental results, theoretical studies have also suggested that convex-shaped proteins are key to forming the cellular protrusions through polymerization of cortical actin ^[24–27]^.

Amongst the I-BAR proteins, IRSp53 has been extensively studied. It links the plasma membrane deformations and associated protrusive activity at the cell boundary to the underlying actin cytoskeleton ^[22]^, by binding to small GTPases, including Cdc42 and Rac1 during cell migration ^[28–32]^. IRSp53 also interacts with negatively charged lipids, including phosphoinositides such as phosphatidylinositol (3,4,5)-trisphosphate and phosphatidylinositol (4,5)-bisphosphate ^[19,33,34]^, which in turn are able to activate Cdc42 and Rac1 through guanine nucleotide exchange factors. The studies on flat 2D surfaces and 3D gels have demonstrated that the depletion of IRSp53 results in impaired protrusive activity at the cell membrane and during cell migration ^[21,22,28,35–37]^. In this study, we investigated if the governing molecular machinery of cellular protrusions and cell migration discovered in the 2D systems and 3D gels extended to suspended 1D fibers. Using suspended fibers of 135 and 500 nm diameter we probed the role of IRSp53 during cell spreading and migration on 1D fibers. Unexpectedly, IRSp53 depletion resulted in reduced actin activity leading to loss of contractility and change in migratory phenotype confirmed by our theoretical model. Overall, our experimental and theoretical study in a biologically relevant 1D fibrous environment describes IRSp53’s unique role in mechano-transduction on 1D fibers, i.e., the actin filament dynamics at the protrusive tip of the elongated cells appears to be essential for the force transduction to the nucleus and for cell migration.

## 2. Results

### 2.1. IRSp53 depletion alters spreading dynamics on suspended fibers but not on flat 2D

In order to quantitate the role of IRSp53 at the interface of membrane dynamics and cytoskeletal contractility, we generated the IRSp53 knockout (KO) U251 glioblastoma cells (**Supplementary Figure S1**) using a CRISPR/Cas9 system, as described previously ^[38]^. We first inquired if IRSp53 depletion caused any effects easily observable under a microscope. We selected a cell-spreading assay composed of suspended fibers of two diameters (high curvature ∼135 nm and low curvature ∼500nm) spaced at least 20 μm apart to achieve spindle-like elongated cell shapes attached to single fibers (**Fig. 1A**). We confirmed the diameters using scanning electron microscopy (SEM) (**Supplementary Figure S2**). Cell spreading experiments on the suspended fibers were timed to capture spreading behavior from a rounded initial state (high circularity) to a more elongated state (low circularity) along fibers (**Fig. 1A, B**). We calculated the area of cells as they spread and found that on the two diameters, both cell types spread at similar rates, and when compared with flat 2D, cells on suspended fibers had smaller areas (**Supplementary Figure S3, Movies M1, and M2**). However, we observed that KO cells took longer times to achieve low circularity values on fibers (longer time constant) than the WT cells on both fiber diameters, while on flat 2D, both cell types remained highly circular during spreading (**Fig. 1C**). Thus, while overall cellular area calculations were not able to distinguish major differences between the two cell types, the time-scale of the circularity metric, driven by protrusive activity on the fibers, was able to clearly and quickly distinguish IRSp53 depletion effects.

**Figure 1:**
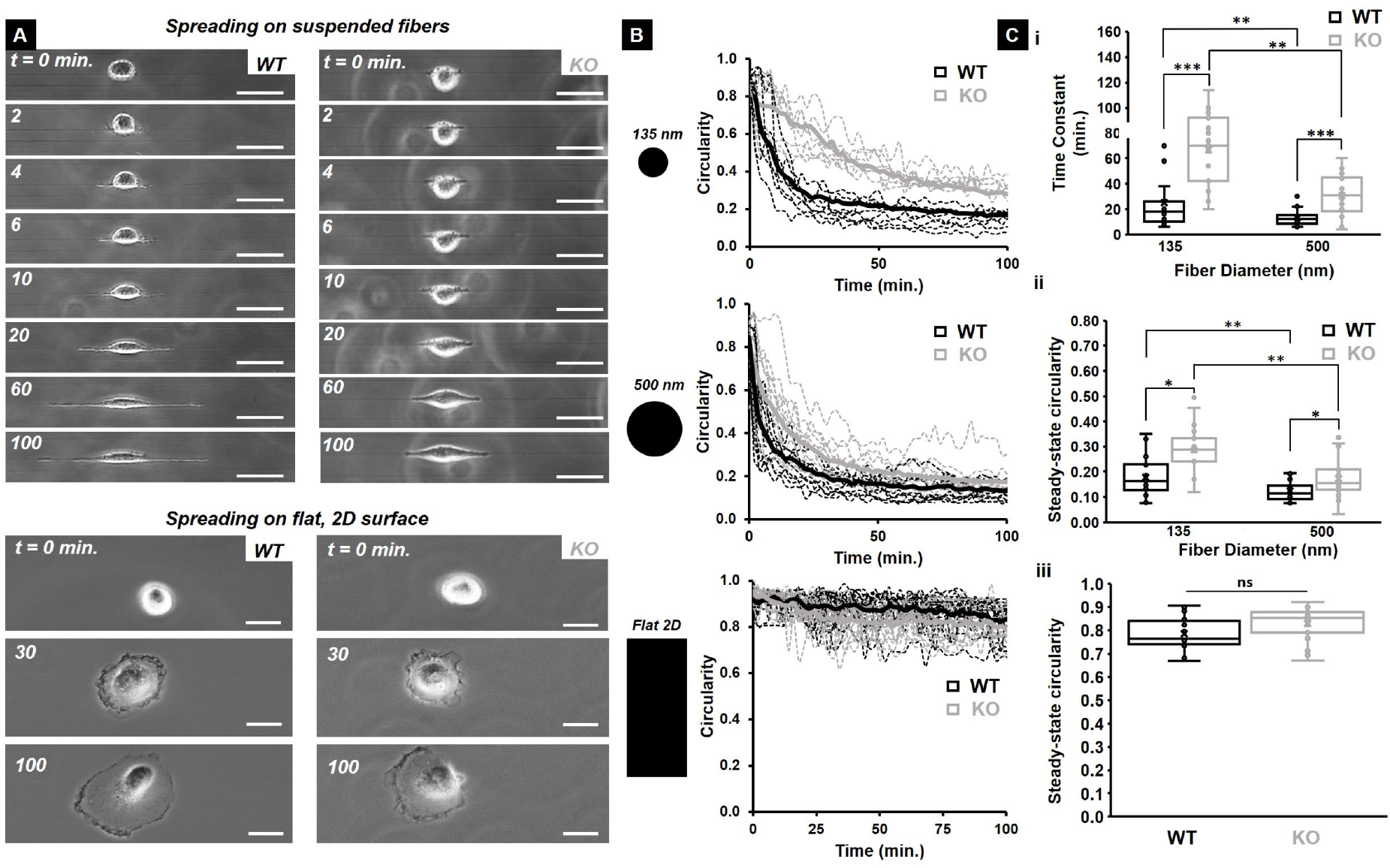
IRSp53 KO cells circularity change is slower compared to WT cells on suspended fibers. (A) Sequence of phase images showing WT cells (left panel) and KO cells (right panel) spreading on 135 nm diameter suspended fibers (top) and on flat, 2D surface (bottom). Scale bars are 20 μm. (B) Circularity profiles for both WT and KO cells on 135 nm and 500 nm diameter suspended fibers and on flat, 2D surface. Black and grey dotted lines represent the individual circularity profiles for WT and KO cells respectively while the solid lines represent average profiles. (C) Quantification of the (i) time constant and (ii) steady state circularity on both 135 nm and 500 nm diameter suspended fibers and (iii) steady state circularity on flat, 2D surface for both WT and KO cells. n =20 for all categories on all substrates. All error bars shown represent standard error of mean.

### 2.2 IRSp53 depletion impairs protrusion dynamics and coiling at the tips of protrusions

Since cell spreading is initiated through the extension of protrusions, and IRSp53 KO cells were taking longer times to achieve lower circularities (elongated shapes) on suspended fibers of both diameters, we inquired if protrusion dynamics were affected by IRSp53 depletion. To quantitate protrusive activity, we used our approach of depositing large diameter (∼2 μm), suspended “base fibers” orthogonal to smaller diameter suspended “protrusive fibers” (**Fig. 2A(i, ii)**) ^[7,8,12]^. As previously reported, this configuration allows bulk cell body migration to be constrained along with the base fiber, while individual protrusive events are elicited and studied along the protrusive fibers. To quantify the protrusion dynamics, we measured the protrusion length (L) and the eccentricity (E, a measure of the protrusion width and shape at its base). Low eccentricity values (E< ∼0.6) signified “rod-like” protrusions, while higher eccentricity values (E> ∼0.8) indicate broader, “kite-shaped” protrusions. Using the combination of the protrusion length and eccentricity allowed us to quantitatively describe a “protrusion cycle.” A typical protrusion cycle commences with the rapid broadening of the protrusion at its base, i.e., an increase in the eccentricity, followed closely by an increase in the protrusion length until the protrusion reaches a maximum length and finally retracts back to the main cell body (**Fig. 2B (i, ii), Movie M3**). We found no significant differences in the averages of maximum protrusion lengths between the KO and WT cells on both fiber diameters tested (**Fig. 2C(i)**). However, we found that on both fiber diameters, the eccentricity (E) was significantly higher for the KO cells suggesting broader protrusions (**Fig. 2C(ii)**). Despite that the maximal lengths were similar, we found that IRSp53-KO cells took a longer time to reach them (**Figure 2C(iii)**), indicating lower protrusive speeds (**Figure 2C(iv)**). Additionally, the KO cells exhibited significant fluctuations (extension and retraction) during their growth phase (**Supplementary Figure S4**).

**Figure 2:**
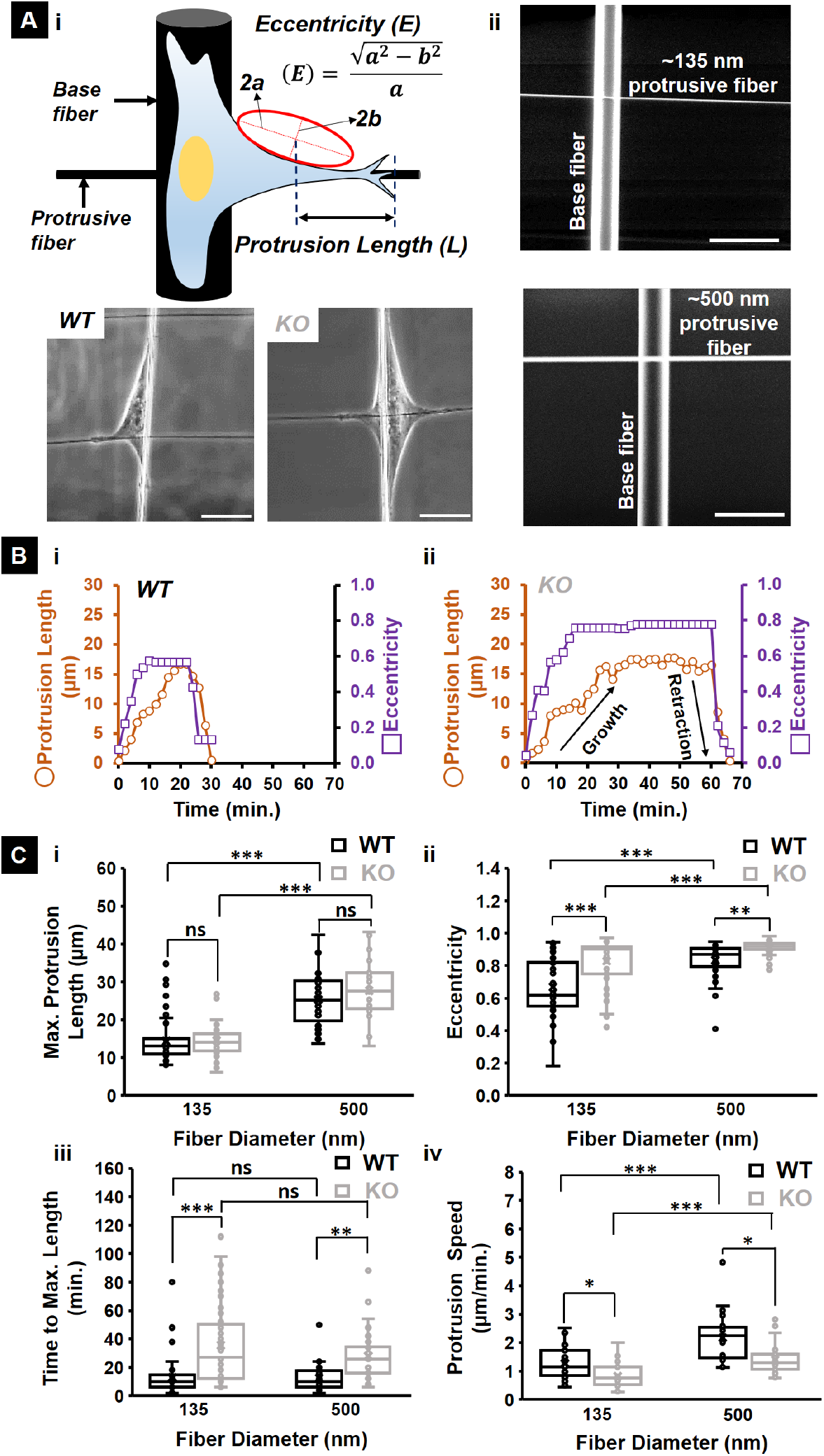
IRSp53 KO glioma cells extend protrusions slower compared to WT cells on suspended fiber networks. (A.i) Schematic showing how the protrusion length and eccentricity are quantified on the fiber networks. (A.ii) Scanning electron microscopy (SEM) images of the fiber networks manufactured using non-electrospinning STEP technique for the protrusion measurements. Scale bars are 5 μm. (A.iii) Brightfield images depicting typical protrusions formed by a WT and KO cell on 135 nm diameter protrusive fibers. Scale bars are 20 μm. Representative protrusive cycles for both WT (B.i) and KO (B.ii) cells highlighting the significant differences in protrusion formation dynamics. (C) Quantifying the differences in (i) maximum protrusion length, (ii) eccentricity at the maximum length, (iii) time taken to reach the maximum length and (iv) protrusion speed between WT and KO cells extending protrusions on both 135 nm and 500 nm diameter protrusive fibers. n values for KO cells are 50 and 38 on 135 nm and 500 nm diameters respectively and for WT cells are 50 and 32 on 135 nm and 500 nm diameters respectively. All error bars shown represent standard error of mean.

Previously, we have demonstrated that cells *coil* (wrap-around the fiber axis) at the protrusion tip (**Fig. 3A(i, ii)**) ^[8,12]^. Based upon our findings of delayed protrusive activity in IRSp53 KO cells, we naturally inquired if these differences translated to differences in the *coiling* cycle occurring at the protrusion tip (**Fig. 3B(i), Movie M4**). While the maximum *coil* width increased with fiber diameter (**Fig. 3B(ii)**), in agreement with our previous findings ^[8]^, the maximum *coil* width was significantly lower for the KO case (**Figure 3B(ii)**). Overall, we found that IRSp53 protein depletion did not impact the protrusion lengths but impaired protrusion dynamics and led to diminished *coil* width at the protrusion tip.

**Figure 3:**
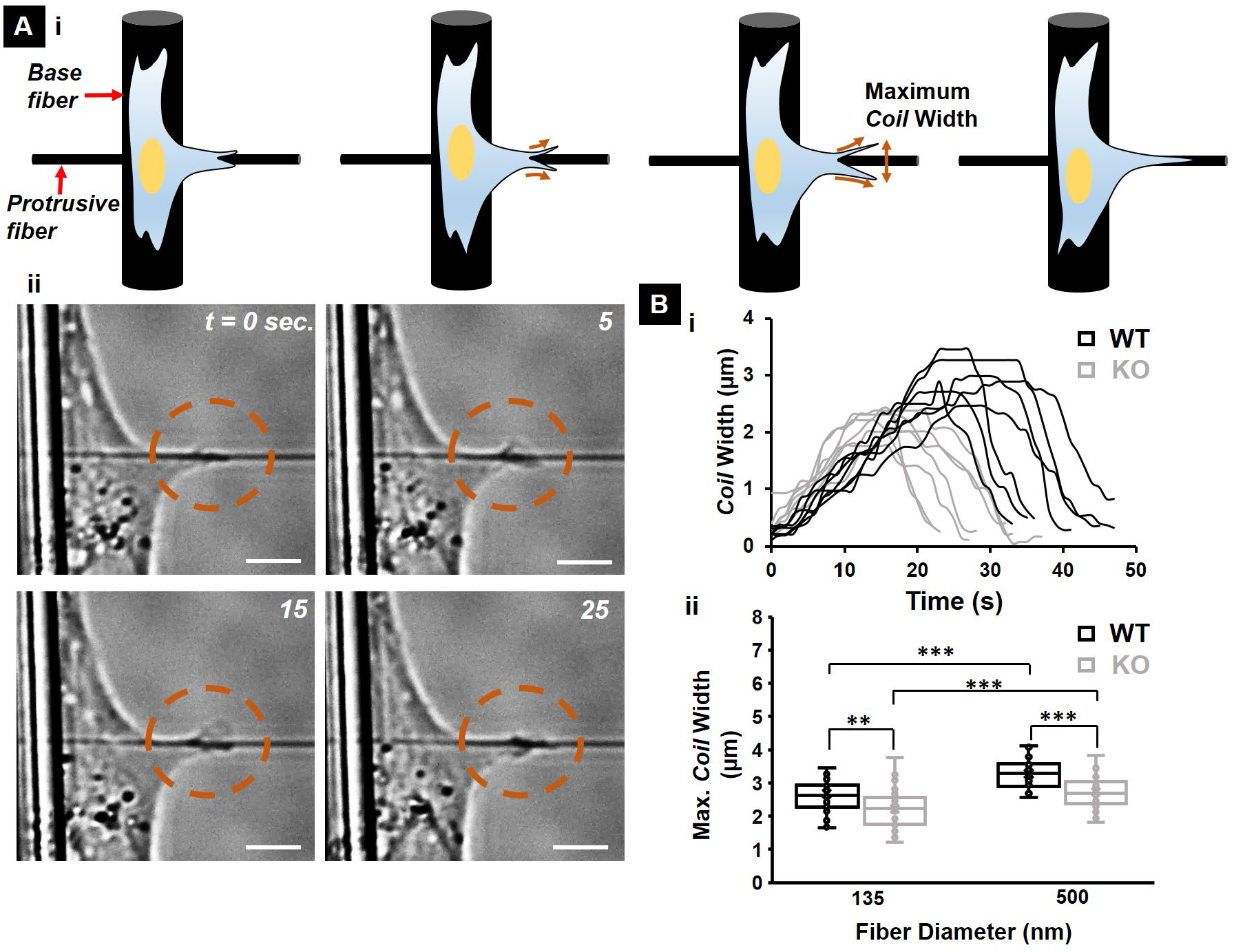
IRSp53 KO cells exhibited lower *coil* widths at the protrusion tip. (A.i) Schematic showing a typical *coiling* cycle with (ii) associated phase images for a KO cell on 135 nm diameter protrusive fiber. Scale bars are 5 μm each. Dashed orange circles highlight the *coiling* in each of the phase images. (B.i) Representative *coiling* cycles for both WT and KO cells on 135 nm diameter protrusive fiber (7 representative profiles for each case). (B.ii) maximum *coil* width highlighting the diminished *coil* width for the KO cells. n = 30 for both WT and KO cells on each of the two fiber diameters investigated. All error bars shown represent standard error of mean.

### 2.3 IRSp53 depletion affects actin networks and contractility

Since delayed cell spreading is associated with a loss of force exertion ^[39]^, we inquired if depletion of IRSp53 resulted in a loss of contractility. We used nanonet force microscopy (NFM) to quantify the forces exerted by single cells ^[11,12,40–42]^, as they spread on two parallel fibers of the same diameters (**Movie M5**). NFM estimates forces by establishing force vectors that originate at focal adhesion clusters (FAC) and are directed along the actin stress fibers (**Fig. 4A**). On suspended fibers, cells form FAC at the poles (**Supplementary Figure S5**; 4 in case of cells spanning two parallel fibers); thus, the overall contractility of the cell estimated over the four FAC is *F*_*cell*_ = ∑ *FAC*. Using fluorescent images of filamentous actin, we first quantified the average stress fiber angle relative to the fiber (*θ*, **Fig. 4 B(i)**) and found no difference between the KO and WT cells on both fiber diameters (**Fig. 4B (ii)**). For both cell types, the average stress fiber angle increased significantly with the increase in the fiber diameter. However, we found that IRSp53 KO cells exerted ∼40% less force than WT cells (**Fig. 4C(i, ii)**). This decrease in contractile force appeared together with a reduction in the density of prominent, cell-length spanning stress fibers in the IRSp53 depleted cells as compared to the WT cells on the two parallel suspended fibers (**Fig. 4D(i)**, and **Supplementary Figure S6**), as well as on a single fiber (**Fig. 4E(i)**). By comparison, the decrease in prominent stress fibers due to the depletion of IRSp53 was not observed on the flat 2D surface (**Supplementary Figure S6, and S7**). In contrast to the stress fiber density, we observed no significant differences in the distribution of focal adhesions in the WT and KO cells, on both the suspended fibers and on a flat 2D surface (**Supplementary Figure S8**).

**Figure 4:**
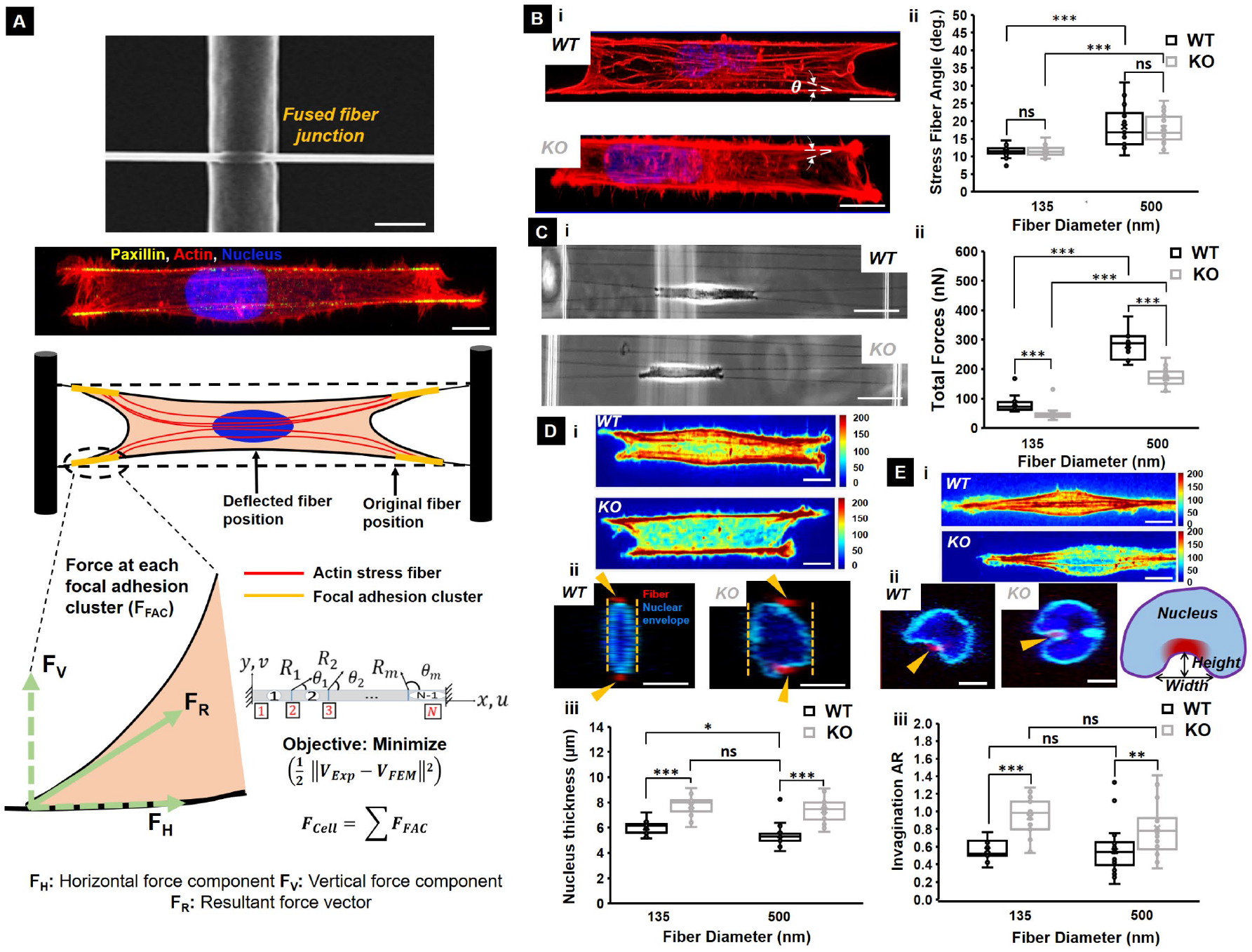
IRSp53 depletion leads to loss of contractility through loss of actin stress fibers. (A) Schematic providing an overview of how forces are calculated using fused nanonets. The SEM image shows a fused fiber junction. Scale bar is 2 μm. The fluorescent image shows actin filaments in red, nucleus in blue and focal adhesion protein paxillin clustering in yellow. Scale bar is 20 μm. NFM establishes force vectors that originate from focal adhesion clusters and are directed along the actin stress fibers. An inverse finite element model minimizes the error between computational and experimental fiber deflections. (B) Representative fluorescence microscopy images of (i) WT (top) and KO (bottom) cells with actin stained in red and (ii) quantification of the stress fiber angles for both cell types on 135 and 500 nm diameter fibers. Scale bars are 10 μm. Dotted white lines in the fluorescent images depict the stress fiber angles. n values are 25 and 25 for the KO cells and 25 and 27 for WT cells on 135 nm and 500 nm diameter fibers respectively. (C) Representative phase images of (i) WT (top) and KO (bottom) cells exerting forces by pulling on suspended fibers with scale bars of 50 μm and (ii) quantification of the forces exerted for both cell types on 135 and 500 nm diameter fibers. n values are 25 for both cell types on each of the two fiber diameters investigated. (D.i) Representative heat maps in arbitrary units of the actin stress fiber distributions for the two cell types with scale bars of 10 μm. (ii) Representative confocal images of WT and KO nucleus cross-section (yz plane) with scale bar of 5 μm, and (iii) quantification of the nucleus thickness. (E.i) Representative heat maps in arbitrary units of the actin stress fiber distributions for the two cell types attached to single fibers in spindle shapes with scale bars of 10 μm, (ii) Representative confocal images of WT and KO nucleus cross-section (yz plane) with scale bar of 5 μm, and (iii) quantification of the nucleus invagination aspect ratio (AR). The schematic in E.ii shows how AR was measured. In the confocal images, the nucleus is in blue, the nuclear envelope is in cyan and the cross-section of the suspended fibers is in red shown using yellow arrowheads. n values are 18 for the nucleus thickness measurements in (D) and 25 for the invagination AR measurements in (E) for both cell types on each of the two fiber diameters investigated. All error bars shown represent standard error of mean.

Given that actomyosin contractility-driven forces have previously been implicated in modulating nuclear deformations ^[43–45]^, we inquired if the diminished contractile forces exerted by the KO cells are manifested in modified nuclear compression. Using confocal microscopy, we measured the nucleus thickness for both KO and WT cells. We found that the KO cells had ∼30% thicker nuclei (indicating reduced nucleus compression, **Figure 4D(ii, iii)**). To confirm that these results were not exclusive to U251, we quantified the stress fiber angles, the exerted forces, and the nucleus thickness for gingival cancer Ca9-22 WT and KO cells (**Supplementary Figure S9**). Motivated by these results on two fiber systems, we enquired if there were any differences in the nucleus shape for cells migrating on single fibers. Confocal imaging revealed invaginations in the nuclear envelope at fiber-specific sites^[46]^. We examined the shape of local invaginations and found a remarkable difference between the WT and KO cells (**Fig. 4E(i, ii)**). The invaginations of the nucleus in the KO cells were sharper and deeper than the invaginations in the WT cells. Quantifying the invagination aspect ratio (height/width) revealed an increase in the invagination aspect ratio for the KO cell nuclei. Altogether, we found that the number density of cell-spanning stress fibers was significantly diminished in the KO cells. Consequently, the KO cells exerted significantly lower forces on the external fibers than the WT cells, which ultimately translated to reduced nuclear compressions and altered invagination shapes at the nucleus-fiber-contact sites.

### 2.4 IRSp53 depletion causes loss of stick-slip migration and a breakdown in nucleus-cytoskeleton coupling

Given that IRSp53 depletion altered force exertion and shape of nuclei, we inquired if these changes ultimately resulted in differences in migration dynamics. Thus, we quantified single-cell migration of both KO and WT cells attached to 500 nm diameter fiber networks as well as on a flat 2D surface. We first characterized the morphology of migrating cells (**Supplementary Figure S10)**, and as expected, both KO and WT cells showed significantly lower circularity and higher aspect ratio on fibers than the flat surface, in agreement with our previous findings ^[47]^. Next, we investigated the migration dynamics and found that WT cells migrated on both substrates using the stick-slip mode (**Movie M6**), whereby in a migration cycle, the leading edge would continue to grow, and the trailing edge would retract in a slingshot manner. In contrast, KO cells exhibited a slower, smooth and sliding migratory behavior with lower persistence (**Fig. 5A(i, ii)**). In the stick-slip mode of migration, WT cells demonstrated synchronous displacement between the bulk cell body and the nucleus, while in KO cells, the nucleus lagged the cell body displacement during migration. To further quantify this behavior, we measured the correlation factor between the nucleus and the cell body displacements, on short time scales of ∼5 minutes. The correlation factor ranges from -1 to 1 (see Methods for details), with values close to 1 indicating highly correlated motion (i.e., the nucleus and cell body move in the same direction together), values close to zero indicating no correlation, and values close to -1 indicating anticorrelated motion. We found that both KO and WT cells on fibers had higher correlation values in general compared to their counterparts on flat 2D indicating a strong coupling between cell body and nucleus during a migration cycle on the 1D fibers. However, depletion of IRSp53 resulted in a loss of correlation on both the fibers and flat substrate, with cells on flat substrate having values closer to zero, indicating almost a complete lack of directional coupling between the nucleus and cell body displacements on short time-scales during migration (**Fig. 5B**). In addition to the reduced coordination during migration, we also found that the KO cells exhibited fewer fluctuations in the cell shape during migration than the WT cells (**Supplementary Figure S11**). Overall, we found that cells moved faster with higher persistence on fibers than on the flat substrates, and that IRSp53 depletion resulted in the loss of stick-slip migratory mode and reduced coupling between the nucleus and the cell body during migration.

**Figure 5:**
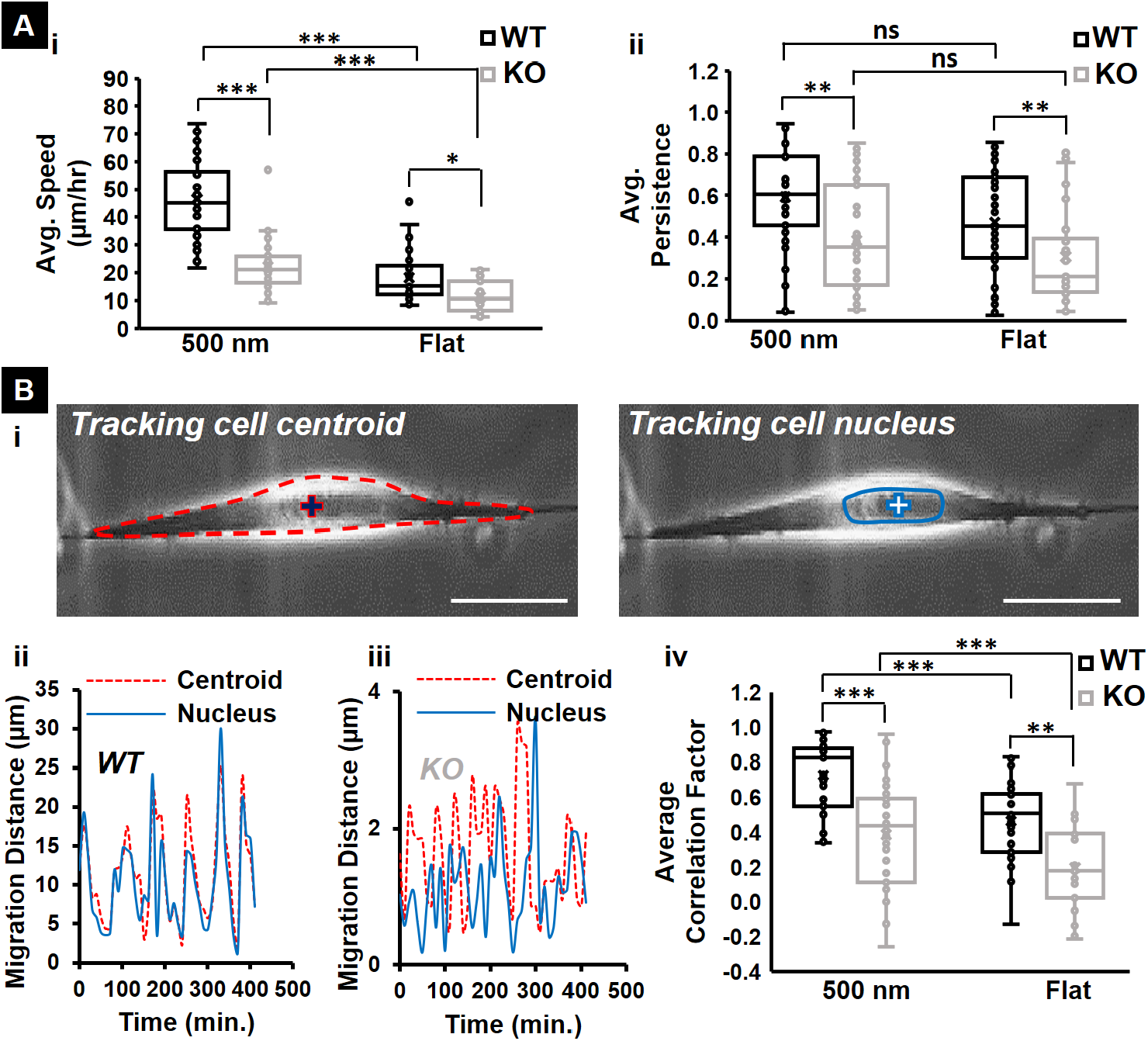
IRSp53 depletion leads to loss of nucleus-cell body coupling and shift in migratory pattern. (A) Quantification of the (i) speed, and (ii) persistence of both WT and KO cells on 500 nm diameter suspended fibers and flat 2D surface (control). (B) Quantifying the synchronicity between the nucleus and cytoskeleton during migration for WT and KO cells. (i) Phase images showing that either the centroid or nucleus can be tracked during migration. Red “+” sign indicates the centroid of the cell body that is outlined by the red dashed boundary. Blue “+” sign indicates the nucleus of the same cell. Typical transient profiles of the distances migrated by the centroid (red) and nucleus (blue) for (ii) WT and (iii) KO cells. (iv) Quantification of the average correlation factor between the centroid and nucleus movement during migration for both WT and KO cells on suspended fibers and flat surface. n values for both cell types are 35 on each substrate. All error bars shown represent standard error of mean.

### 2.5 Restoration of IRSp53 recovered WT behavior

Next, we confirmed that the impaired *coiling* dynamics at the protrusion tip, reduced contractility, and hindered migration dynamics were due to depletion of IRSp53 by quantifying the effects of restoring IRSp53. We expressed IRSp53 in the IRSp53 depleted cells (KO+IRSp53 cell line) and observed that these cells had a protrusive behavior similar to those of WT cells (**Fig. 6A**). Furthermore, KO+IRSp53 cells exerted similar forces, resulting in increased nuclear compression and recovery of nucleus thickness similar to WT cells (**Fig. 6B**). Finally, reconstituted cells recovered stick-slip migration dynamics and nuclear invagination aspect ratios similar to that of the WT cells (**Fig. 6C**). Altogether, we found that IRSp53 KO cells reconstituted with IRSp53 protein (KO+IRSp53 cell line) recovered WT functionalities.

**Figure 6:**
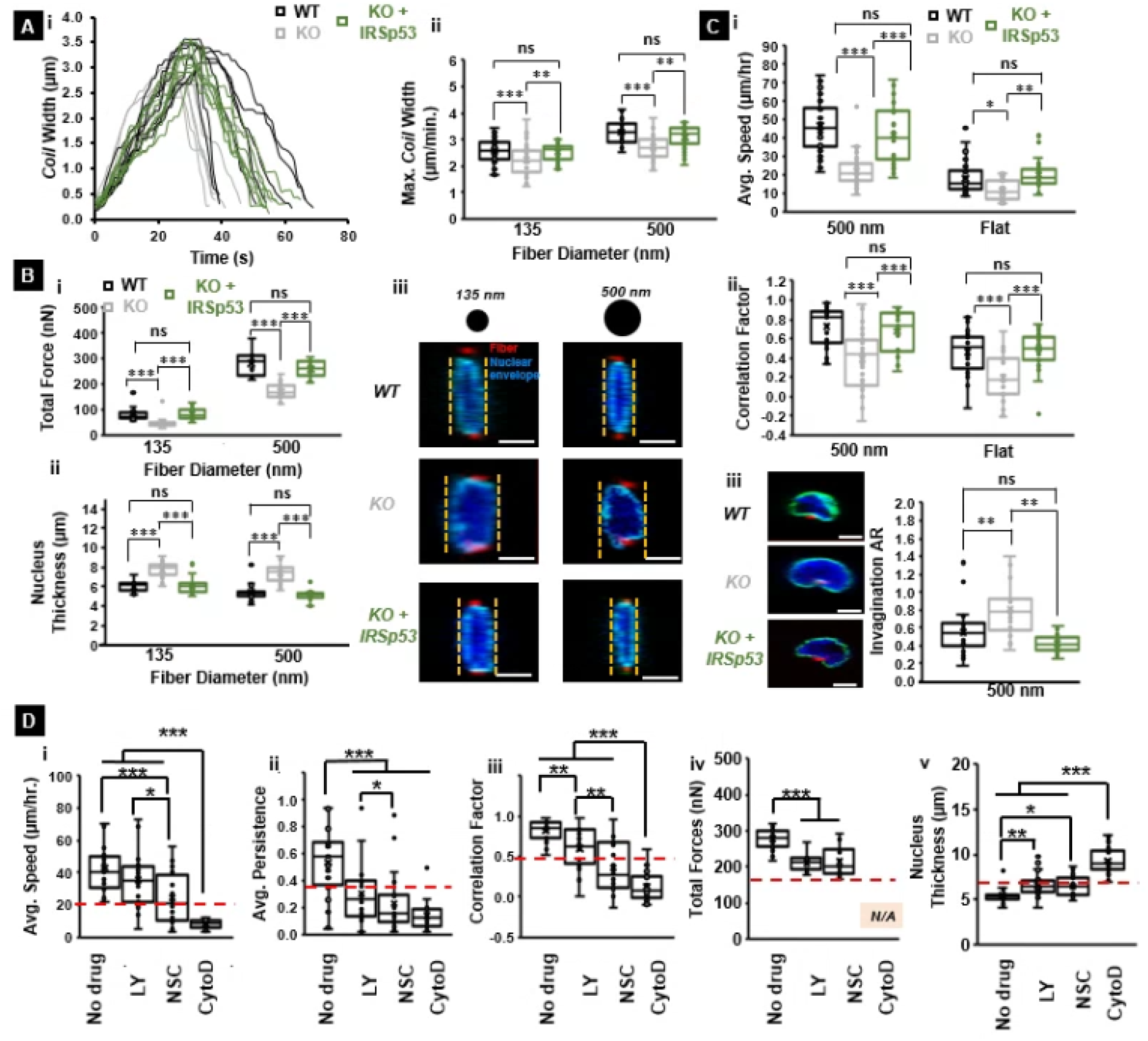
IRSp53 reconstitution in KO cells recovers WT cell function. (A.i) Representative coiling cycle profiles for IRSp53 KO, WT and IRSp53 reconstituted cells on 500 nm diameter suspended fibers. 8 representative profiles selected for each cell category. Quantification of (A.ii) maximum coil width for all three cell types on both 135 nm and 500 nm diameter protrusive fibers. n values are 30 for KO cells, 30 for WT cells and 35 for IRSp53 reconstituted cells on each of the two fiber diameters tested. (B) Quantification of the (i) total forces exerted and (ii) nucleus thickness of IRSp53 KO, WT and IRSp53 reconstituted cells on both 135 nm and 500 nm diameter fibers. (B.iii) Representative confocal images showing that reconstitution of IRSp53 in IRSp53 deficient cells leads to recovery of nucleus thickness similar to the WT cells. n values for the force calculations are 25 for all cell categories on both fiber diameters tested. n values for the nucleus thickness measurements are 18 for the KO cells on both fiber diameters, 18 for the WT cells on both fiber diameters, 18 and 22 for the IRSp53 reconstituted cells on the 135 nm and 500 nm diameter fibers respectively. In the confocal images, the nucleus is in blue, the nuclear envelope is in cyan and the cross-section of the suspended fibers is in red. Scale bars are 5 μm.

### 2.6 Role of PI-3 kinase and Rac in 1D migration and nuclear shape

To examine the signaling cascade involved in the coupling between the nucleus and the cellular protrusions on 1D fibers, we treated the WT and KO cells with the PI-3 kinase inhibitor LY294002 and Rac1 inhibitor NSC23766, as well as the actin polymerization inhibitor cytochalasin D (**Fig. 6D**). The cell migration speed of both WT and KO cells, as determined by the movement of centroid of the cells (as in **Fig.1**), was decreased by treatment of cells with LY294002 and NSC23766 (**Supplementary Figure S12** for KO cells). While KO cells had reduced migration persistence, the persistence of KO cells was further decreased by NSC23766 treatments. WT cells exhibited decreased persistence when treated with LY294002 and NSC23766. Treatment with Cytochalasin D of both cells abrogated the migratory response (**Fig. 6D (i, ii)**). Furthermore, the correlation of the movement of the nucleus and that of the cell centroid in the WT cells was uncoupled by the LY294002 and NSC23766 treatments (**Fig. 6D(iii)**). However, the correlation of the KO cells was not further reduced by treatment with LY294002 treatment but was reduced with NSC23766 treatment, presumably because IRSp53 mediated the PI-3 kinase signaling but did not mediate the Rac1 signaling.

We also measured the total force and nucleus thickness in the presence of these drugs and found a considerable decrease of the force and a corresponding increase of the nuclear thickness with the drug treatments (**Fig. 6D**). Furthermore, the treatment of the cells with the inhibitor of the actin filament turnover, cytochalasin D, resulted in the complete loss of speed, persistence, correlation, and force, as well as the increase of the nuclear thickness (**Fig. 6D(iv, v)**). These results suggest that PI-3 kinase and Rac1 couple 1D migration and nuclear shape. Thus, IRSp53 plays an active role downstream of PI 3-kinase.

### 2.7 Theoretical analysis of IRSp53 effects in 1D migration

We then inquired if we could describe the migratory behavior of IRSp53 KO cells using a theoretical model of one-dimensional cell migration ^[4]^. The model is coarse-grained and simplified, which allows us to qualitatively relate the observed changes in migration characteristics to possible effects of IRSp53 on the microscopic parameters of the cell migration mechanism robustly. One-dimensional elongated cells have actin polymerization activity within this model at their two opposing ends (**Fig. 7A**), where the adhesion to the substrate is also localized (Hennig et al., 2020). The net actin retrograde flow in the cell, given by the difference between the flows from both ends, adverts polarity cues that transiently diffuse or adsorb to the treadmilling actin ^[48]^. For the full theoretical description, we refer the reader to the methods section. In the model, the cell length plays a crucial role: when the cell length is smaller than a critical value *l*_*c*_, the actin flow can not form a large gradient of the polarity cue, and the small difference along the cell length is not sufficient to maintain a polarized cell with a finite net flow. In this case, the actin polymerization activity drives symmetric cell elongation at both ends of the cell until the cell reaches a steady-state length *l*_*p*_, where the cell’s elasticity balances the protrusive forces. We plot the phase diagram of the different migration patterns of the cell, as predicted by our model (**Fig. 7B**) ^[4]^. In the phase diagram, we find that a non-motile phase (*l*_*p*_ < *l*_*c*_) occurs when either the cell-substrate adhesion is low (denoted by the parameter *r*), or when the maximal actin retrograde speed (denoted by *β*) is low. In both conditions, the polymerization of actin at the cell’s ends does not generate protrusive forces that are sufficiently large in order to elongate the cell above the critical length *l*_*c*_, which is needed in order to polarize (**Fig. 7C(i-iii)**). Beyond the non-motile regime, for larger *r* or *β*, we predict a regime where cells migrate smoothly (**Fig. 7D**), and as *β* increases further, the cell is predicted to transition into stick-slip migration mode (**Fig. 7E, F).** Comparing the theoretical model to the experimental observations of decreased protrusive speed and low force exertion, we can therefore propose that the IRSp53-KO cells have a lower maximal actin treadmilling speed (*β*) which places them in the slower and smooth migration or non-motile regimes (points i, ii in Fig. 7B, compare to experimental data in **Fig. 7G**), while the WT cells have a high level of *β* at their edges, placing them in the stick-slip migration regime (points iii, iv in Fig. 7B and shown with experimental data in **Fig. 7H**)

**Figure 7:**
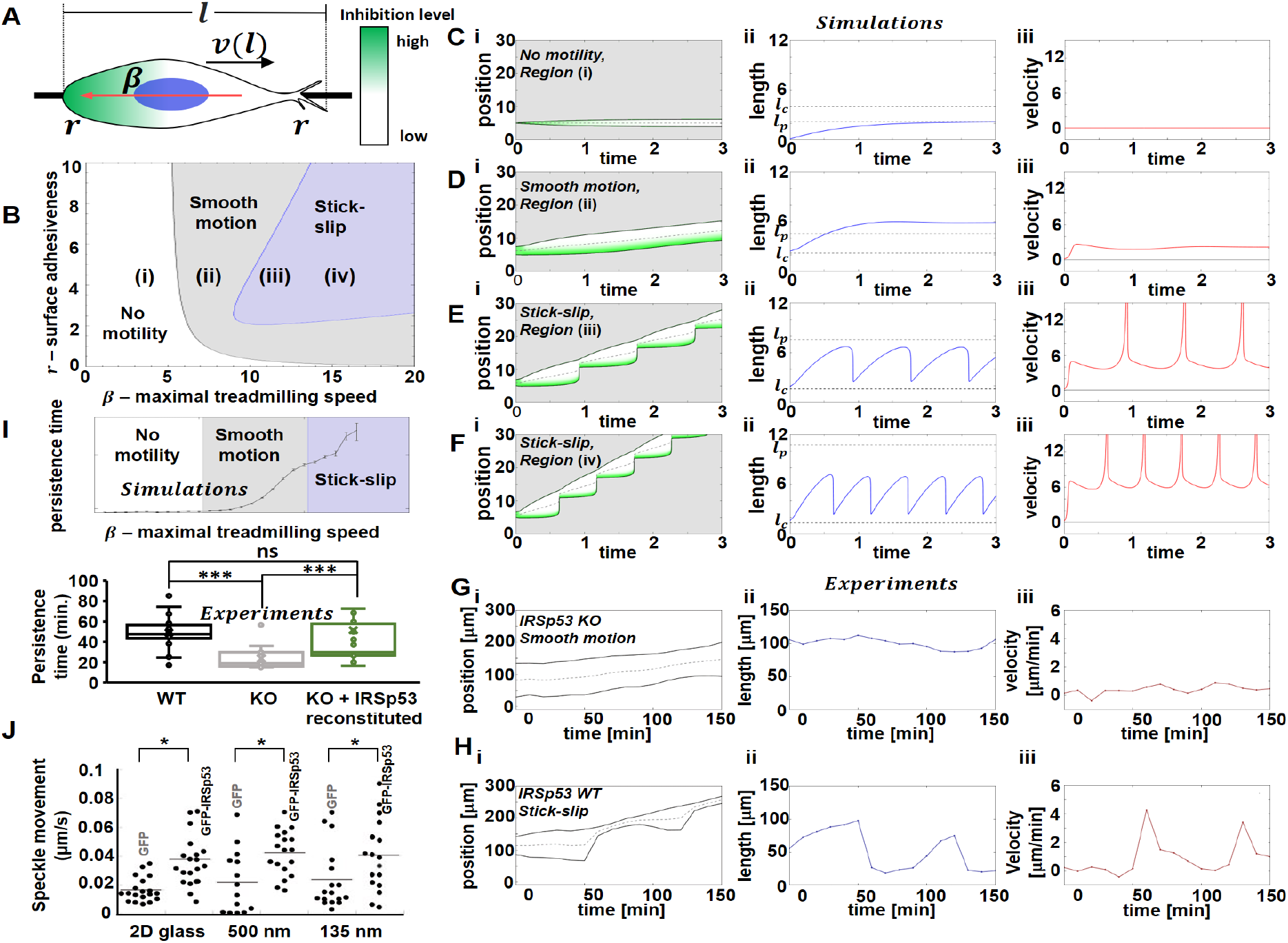
Relating the migration patterns of the IRSp53 KO and WT cells on suspended fibers to a theoretical model of one-dimensional cell migration. (A) Illustration of the theoretical model. A cell of length *l* migrating on a linear 1D fiber. *v*(*l*) represents the velocity of the cell’s center of mass, which depends on the cell length. *β* represents the maximal actin treadmilling flow velocity (red arrow), driven by a gradient of an inhibitory polarity cue (depicted by the colors green/white for high/low inhibition). *r* represents the local cell-surface adhesiveness at the cell edges, and the blue circle denotes the location of the nucleus. (B) A theoretical phase-diagram denoting the different migration patterns as function of the adhesiveness of the substrate (*r*) and the maximal actin treadmilling flow (*β*). Fixing the adhesion value, with increasing *β* the model predicts: (i) cells that are non-motile, elongating symmetrically (up to a maximal length *l*_*p*_) but do not cross the critical length for polarization *l*_*c*_ shown in (C (i-iii)), (ii) migrating smoothly with speed that increases with *β* shown in (D (i-iii)), (iii- iv) exhibiting stick-slip migration, with speed and stick-slip frequency increasing with *β* shown in (E (i-iii) and F (i-iii)). For each condition we plot: (i) The kymographs, where the green/white color gradient is the polarity cue gradient (high/low) and the gray dashed line is the position of the center of mass, (ii) The length time series in blue, and *l*_*p*_/*l*_*c*_ in dashed gray/black line, and (iii) the center-of-mass velocity in red. (G,H) The theoretical results are compared to experiments of IRSp53 KO (G) and WT (H) cells: (i) typical kymographs where the dashed gray line is the nucleus position, (ii) cell length, and (iii) nucleus velocity time series. (I) Top panel: simulation results for the persistence time as a function of the strength of the maximal actin treadmilling flow (*β*). The parameters used for the calculations of the model are presented in the methods section. Bottom panel: experimental measurements shows IRSp53 reconstituted KO cells recover WT persistence times (n=35 per category for cells on suspended fibers and 30 per category for cells on flat). (J) Speckle movement, where each dot represents a measure of the actin retrograde flow velocity at the leading edge of a IRSp53-KO cell expressing GFP and IRSp53-KO cell with GFP- IRSp53.

The model, thus, naturally explains the increased persistence in the WT cells compared to the IRSp53-KO cells (Fig. 5A(ii)). IRSp53 is known to couple with the actin polymerization machinery and we therefore propose to correlate it with the parameter *β*. With the increase of *β*, the speed of the retrograde actin flow increases, which in turn gives rise to a larger front-back gradient of the polarity cue. Thus, cells with a higher speed of the internal actin flow have a higher persistent time, remaining polarized along one direction of motion (in 1D), despite various internal noise sources; a prominent result of the Universal Coupling of Speed and Persistence (UCSP) model ^[4,48]^. Indeed, our model predictions match the decreased persistence times observed for IRSp53-KO compared to WT cells (**Fig. 7I**).

We tested our prediction of WT cells having high actin polymerization at the leading edge by analyzing the retrograde flow of actin using the Halo-tagged actin introduced into the WT and KO cells. Using speckle microscopy, we generated actin flow kymographs correlating with the extent of the actin polymerization at the leading edge (Supplementary Figure S13) ^[49]^. We found that the retrograde actin flow was faster in the IRSp53-KO cells expressing IRSp53 than in the IRSp53-KO cells expressing GFP, both for the 2D flat surface and the fibers (Fig. 7J). Overall, our theoretical 1D model of cell migration describes the transition of slip-stick migratory behavior to a smooth low-speed migratory mode in cells depleted of IRSp53 through a loss of actin polymerization at the cell’s leading edges, which we confirmed experimentally.

## 3. Discussion

In this study, we examined the molecular mechanism of the 1D cell migration. Because of the characteristic protrusions of the cells on the 1D fibers, we focused on IRSp53 that links the membrane to the actin cytoskeleton in the protrusion in 2D or 3D environments. We established a role of IRSp53 in mediating protrusive activity on the 1D fibers, where cells are forced to adapt to the elongated shape and migrate in “stick-slip” migration in U-251 glioma cells and Ca9-22 cells. IRSp53 was found to be essential for force generation during the 1D migration, presumably through the actin filament formation at the edges of the cellular protrusions. IRSp53-related actin activity at the cell edges is linked to the formation of cell-spanning stress fibers that strongly affect nuclear shape and movement coupling during migration.

The studies on cell migration and IRSp53 have previously been mostly conducted on flat 2D substrates. During 2D and 3D cell migration, cellular protrusions occur by the reorganization of the actin cytoskeleton, which is under the control of small GTPases, including Cdc42 and Rac1. IRSp53 can transmit the signals from both small GTPases to the membrane deformation and presumably to the actin filament dynamics through WASP family proteins ^[31,37]^. Importantly, the membrane recruitment of IRSp53, WASP family proteins, as well as the activation of the small GTPases, are under the control of the phosphoinositide metabolisms, of which the activation of PI-3 kinase is essential ^[19]^. Our pharmacological studies indicated that the protrusive structure formation was dependent on Rac1 small GTPase, PI-3 kinase, and actin polymerization. Interestingly, PI-3 kinase dependency was absent and Rac1 dependency was active in IRSp53 KO cells. Therefore, IRSp53 was suggested to be under PI-3 kinase, but not Rac1, indicating the IRSp53 was involved downstream of Cdc42, which will be studied in the future.

In our study, IRSp53 depletion resulted in hindered migration dynamics in both cells migrating on 1D fibers and flat surfaces; however, the reduction in both speed and persistence of migration was more significant on the fibers (**Fig.5**) suggesting that the fiber networks have greater sensitivity in teasing out such differences. The 1D fibers force the cells to be elongated, where the protrusive structure would naturally occur at the tips of the cells. In such an elongated shape in the 1D environment, actin stress-fibers that span from the tips of the cells to the nucleus play an important role in determining the nuclear shape and in mediating the coupling of the nuclear movements to those of the plasma membrane. Our observation that loss of IRSp53 leads to a slower and smoother mode of migration in which the cell body and nucleus movement are not well correlated could perhaps be due to a mechanical constraint at sites of invagination on 1D fibers.

Although our theoretical framework can explain the decoupling of the nucleus movement by the decrease of the actin filament turnover at the tips, still, IRSp53 might be involved in the molecular coupling of actin filaments to the nucleus. There are abundant IRSp53 in the cytoplasm, and thus it is still possible that IRSp53 may play an unexpected role on the nucleus, as well as in mediating the actin filament reorganization and linking the membrane to the actin filaments at the cellular protrusions. There are several linker molecules between the cytoplasm and nucleus. The Linker of Nucleoskeleton and Cytoskeleton (LINC) complex plays a key role in facilitating connections between the nucleus and cytoplasm ^[50]^. Specifically, the Nesprin family of proteins in the LINC complex connects the actin filaments, microtubules, and intermediate filaments in the cytoplasm to the SUN complex proteins, which in turn are linked to the nuclear envelope ^[51,52]^. Thus, it is tempting to speculate that knocking down the IRSp53 protein could potentially disrupt the pathway through which the LINC complex connects the cytoplasm and nucleus, resulting in asynchronous migratory behavior.

## 4. Conclusion

In conclusion, our use of ECM-mimicking suspended 1D fibers provide insights into cell-fiber interactions. Using fibers of varying diameters shows the sensitivity of cell contractility to the curvature and reduced dimensionality, that is more difficult to expose clearly on flat 2D substrates. We uncover the consequences of IRSp53 depletion for the cytoskeleton organization on 1D fibers, and resulting effects on the traction forces exerted by the cells and their mode of migration. Altogether, we provide new insights into the role of IRSp53 in mechanotransduction during 1D spreading and 1D/3D migration of cells.

## 5. Experimental section/Methods

### 5.1 Fiber Network Fabrication

The previously reported non-electrospinning Spinneret Based Tunable Engineered Parameters (STEP) method was used to fabricate all the suspended fiber networks used in this study. Briefly, polystyrene (PS, Scientific Polymer Products, Ontario, NY) of ∼2×10^6^ g/mol molecular weight was dissolved in xylene (Thermo Fischer Scientific, Pittsburgh, PA) at 6% (w/w) concentration and 10% (w/w) concentration to prepare the solutions for spinning the ∼135 nm and ∼500 nm diameter fibers respectively. To spin the ∼2 μm base fibers for the protrusion, *coiling*, and force studies, a 5% (w/w) concentration solution of polystyrene of ∼15×10^6^ g/mol molecular weight was used. The solutions were prepared at least two weeks prior to spinning the fiber networks.

### 5.2 Scanning Electron Microscopy

Environmental Scanning Electron Microscope (ESEM) was used to take images of the suspended fibers in order to confirm the fiber diameter. Prior to imaging the scaffolds, they were coated with a 7 nm thick layer of Platinum-Palladium using a Leica sputter coater (Leica, Wetzlar, Germany). The images were taken at an electron beam voltage of 10 kV and a spot size of 3.5 using the Everhart-Thornley detector (ETD). The working distance was maintained at ∼11 mm. An appropriate magnification factor was used depending on the application.

### 5.3 Cell Culture and drug studies

U-251 and Ca9-22 cells were obtained from the Japanese Collection of Research Bioresources Cell Bank. The IRSp53 knockout (KO) cells were generated by the CRISPR/Cas9 system, as described previously ^[38]^. The guide RNA targeting the first exon of IRSp53 (CCATGGCGATGAAGTTCCGG) was designed using the server http://crispr.mit.edu and inserted into the pX330 vector ^[38]^. After transfection, the cells were cloned by monitoring the GFP fluorescence from the reporter plasmid pCAG-EGxxFP with the IRSp53 genome fragment using a fluorescence-activated cell sorter [FACSAria (BD)] ^[53]^. The expression of GFP or GFP-IRSp53 in the IRSp53 knockout cells was performed by the retrovirus-mediated gene transfer, as described previously ^[53]^. All cell lines were cultured in high glucose DMEM (Thermo Fisher Scientific) supplemented with 10% bovine calf serum (Thermo Fischer Scientific) and 1% penicillin-streptomycin solution (Thermo Fischer Scientific) and stored in an incubator at 37°C in 5% CO_2_ and humidified conditions. For the pharmacological inhibitor studies, cells were initially seeded as described previously on the nanofiber scaffolds. Subsequently, we added either 20 μm LY294002 (Millipore Sigma, St. Louis, Missouri for PI-3 kinase), 75 μm NSC23766 (Millipore Sigma for Rac1), or 2 μm of Cytochalasin D (Millipore Sigma for Actin) to the cell culture media. The cells were incubated for 2 hours. Following the incubation period, the cells were imaged as previously described.

### 5.4 Cell Seeding and Experiment

In preparation for the experiments, the scaffolds were first affixed to the glass bottom of 6-well dishes (MatTek Corp., Ashland, MA) using sterile, high-vacuum grease (Dow Corning, Midland, MI). The scaffolds were then soaked in 70% ethanol for disinfection, followed by two phosphate-buffered saline (PBS) washes (Thermo Fisher Scientific). Subsequently, the fibers were coated with either 4 μg/ml fibronectin (Invitrogen, Carsbad, CA) or 4 μg/ml rhodamine fibronectin (Cytoskeleton Inc., Denver, CO) for 2 hours prior to cell seeding to aid cell attachment to the fibers. Once the cell culture reached ∼80% confluency, 0.25% Trypsin (ATCC, Manassas, VA) was added, and the culture was incubated for ∼ 5 minutes. After the cells detached from the flask surface, 3 ml of fresh cell media was added to dilute the effect of the trypsin. The entire solution was then placed in a centrifuge at 1000 RPM for 5 minutes. Following the centrifugation, the media was aspirated, and the cells were resuspended in fresh media. Finally, cells were seeded at a density of ∼3000,000 cells/ml on the scaffolds.

### 5.5 Immunostaining

Cells were fixed in 4% paraformaldehyde (Santa Cruz Biotechnology, Dallas, Texas), dissolved in PBS for 15 minutes, and rinsed in PBS twice. 300 μl of permeabilization solution (0.1% Triton-X-100 in PBS) was then added to permeabilize the cells. After 15 minutes, the permeabilization solution was aspirated, and the cells were blocked using 10% goat serum in PBS for 30 minutes. Following this, the cells were incubated with either the anti-lamin A/C primary antibody or the anti-paxillin antibody (Abcam, Cambridge, United Kingdom) dissolved in antibody dilution buffer (0.3% Triton-X-100 and 1% BSA in PBS) at a ratio of either 1:200 (for lamin) or 1:500 (for paxillin) for either 2 hours at 37°C (for lamin) or 16 hours at 4°C (for paxillin). After the incubation period, the secondary antibody Alexa Fluor 647 goat anti-mouse (Invitrogen, Carlsbad, CA) was diluted in the antibody dilution buffer at the ratio of 1:500 and added to the wells for lamin. For paxillin, the secondary antibody Alexa Fluor 488 goat anti-rabbit (Invitrogen) was diluted in the antibody dilution buffer at a ratio of 1:350 and added to the wells. For imaging the filamentous actin for the stress fiber angle measurements, no primary antibody was added to the well. In this case, after permeabilization, rhodamine-phalloidin (Abcam) was dissolved in the antibody dilution buffer at the ratio of 1:80 and added directly to the well. After immunochemistry, the samples were stored in a dark place for 45 minutes, followed by 3 PBS washes. Finally, the nuclei were counterstained with 300 nM of DAPI (Invitrogen) for 15 minutes. The scaffolds were kept hydrated in 2 ml of PBS and imaged using a 63× (water-based immersion) magnification.

### 5.6 Microscopy and Imaging

The cells were imaged using the AxioObserver Z.1 (with mRm camera) microscope (Carl Zeiss, Germany) at 20× for the protrusion, spreading, force, and migration studies and at 63× (water immersion objective) magnification for the *coiling* studies. The imaging intervals used were 5 minutes for the migration study, 3 minutes for the force and spreading studies, 2 minutes for the protrusion study, and 1 second for the *coiling* study. All the videos were analyzed using ImageJ (National Institutes of Health, Bethesda, MD).

For confocal microscopy of the immunostained samples, the cells were imaged using the LSM 880 confocal microscope (Carl Zeiss, Germany) at 63× (water immersion objective). The slice thickness was set to 0.36 μm for all the images, and appropriate laser powers were selected for the different lasers.

### 5.7 Analysis of Biophysical Metrics

For the protrusion analysis, the maximum protrusion length was calculated as described previously. Briefly, the distance from the base fiber to the protrusion tip was first measured (*L*_*base*_). Subsequently, the largest possible ellipse was fit along the curvature of the protrusion such that one end of the ellipse coincided with a point on the protrusion that was at a distance of 0.8 × *L*_*base*_ from the base fiber. Finally, the protrusion length (*L*) was measured as the distance from the tip of the protrusion to the projection of the intersection of the major and minor axes of this ellipse with the protrusive fiber. The eccentricity of the protrusion (*E*) was calculated as follows:

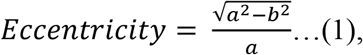

where *a* and *b* are half of the length of the major and minor axis, respectively of the ellipse fit to the protrusion (**Supplementary Figure S14**). The time taken to reach the maximum protrusion length was calculated as the total time taken from the first instance that a protrusion reached a threshold minimum length of 5 μm (this threshold was established in order to discount short-lived membrane spikes and blebs) till the protrusion reached the maximum length. To calculate the average protrusion speed, the protrusion length was recorded for every frame (2 minutes). The instantaneous protrusion speed was first calculated as follows (in μm/hr):

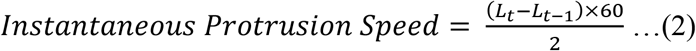

where *L*_*t*−1_and *L*_*t*_ is the protrusion length at any given frame and *L*_*t*−1_ is the protrusion length in the previous frame. Finally, the average protrusion speed was calculated as the average of all the instantaneous protrusion speeds. To determine the percentage slope changes in the protrusion length during a protrusion cycle, the number of times the slope of the protrusion cycle changed sign from positive to negative was first recorded. This was then divided by the total number of time points in the protrusion cycle.

The *coiling* dynamics at the tip of the protrusion were calculated as previously described ^[8]^. Briefly, the maximum *coil* width was calculated as the largest *coil* width during a *coiling* cycle

For the spreading analysis, the circularity of the cell was defined as follows:

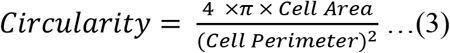

The value for circularity ranges from 0-1 wherein a value closer to 1 indicates a circular shape while a value closer to 0 indicates a more “straight-line” shape. The steady-state circularity was determined at ninety minutes after the initial seeding of the cells as there was no significant change in the circularity beyond this time point. The time constant for each circularity profile was calculated from the exponential decay equation shown below:

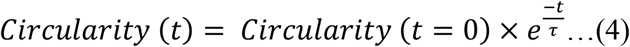

where *τ* represents the time constant and is calculated as the time at which:

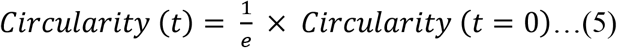

In order to generate the actin heat maps, fluorescent images of the actin stress fibers were first converted to grayscale using ImageJ. Subsequently, the greyscale images were imported into MATLAB and the function “colormap” was used to generate heat maps. For the focal adhesion cluster (FAC) analysis, the FAC length was measured as the longest, continuous length of paxillin clustering. The FAC distance from the nucleus was measured to be the distance from the centroid of the nucleus to the starting point of the FAC length along the suspended fiber axis. The invagination ratio for cells suspended on single fibers was quantified from the yz nucleus mid-plane projection obtained from confocal microscopy imaging as shown below:

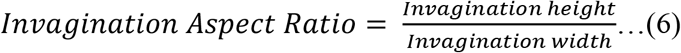

For the migration analysis, cells were manually tracked using ImageJ and the x,y location of the cell centroid was recorded for every second frame (*i.e*., every ten minutes). The instantaneous speed was then calculated as follows (in μm/hr):

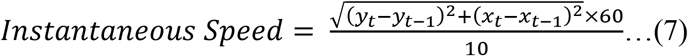

where (*x*_*t*−1_, *y*_*t*−1_) are the coordinates of the centroid of a cell at any given frame in μm while (*x*_*t*_, *y*_*t*_) are the coordinates of the centroid of the same cell three frames (*i.e*., twelve minutes) later. The overall average speed of the cell was then calculated as the average of all the instantaneous speed values. The persistence of migration was calculated as follows:

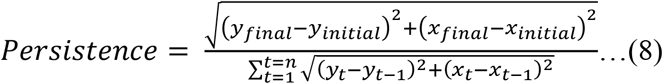

where (*x*_*final*_, *y*_*final*_) are the coordinates of the centroid of a cell at the last frame (n^th^ frame) tracked in μm while (*x*_*initial*_, *y*_*initial*_) are the coordinates of the centroid of the same cell at the first frame tracked. The denominator is defined similar as above for the instantaneous speed. The correlation factor used for determining nucleus-centroid coupling during migration was calculated as follows:

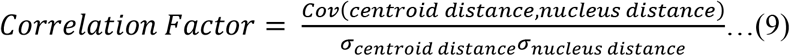

where *cov*(*centroid distance nucleus distance*) represents the covariance of the centroid distance and nucleus distance and *σ* represents the standard deviation. The Shapiro-Wilks normality test was used to ensure the normal distribution of data before calculating the correlation factor. To determine the average change in the area during the migration period, the magnitude of the difference in the area was calculated between every two frames (i.e., 10 minutes), and the average of all these differences was taken. A similar approach was followed to calculate the average changes in both perimeter and circularity.

### 5.8 Force Model for Nanonet Force Microscopy

In order to quantify the forces from the fiber deflections, the fiber deflection was tracked for three randomly selected, consecutive frames and was analyzed in MATLAB (2017a) using our previously reported methods ^[11,42,54]^. Briefly, the ∼135 nm or ∼500 nm diameter horizontal force fibers were modeled as beams with fixed-fixed boundary conditions as they were fused to larger diameter, strut-like, vertical base fibers at either end. Force vectors were established at FAC clusters at the poles and directed along the actin stress fibers. A custom finite element model was used to obtain the force fiber deflection profile based on an arbitrary initial force input. Subsequently, the error between the fiber profile predicted by the model and the experimentally tracked fiber profile was minimized using an optimization frame while simultaneously updating the force values iteratively. The average force was finally calculated as the average of the three consecutive frames selected.

### 5.9 Theoretical Model of 1D cell migration

To describe theoretically the migration of a single U251 glioblastoma cell on a linear fiber, the cell was modeled as a dynamic system (Figure S10) and described by the following equations ^[4]^:

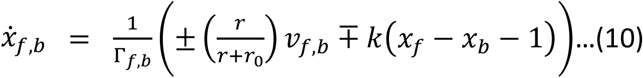

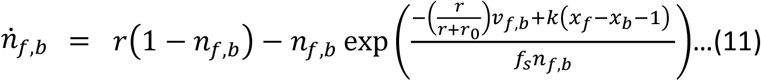

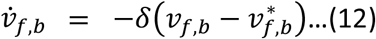

where the state variables 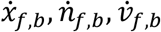 represent the position, adhesion concentration, and actin polymerization speed at the front and back edges of the one-dimensional cell, respectively. The parameter *r* represents the cell-surface adhesiveness due to the binding and unbinding of adhesions, and the parameter *r*_0_ represents an effective saturated cell-surface adhesiveness. The parameter *k* represents the cell elasticity (mean cell spring constant). The parameters *f*_*d*_ and *κ* are associated with the mechanical properties of the adhesions^[4]^. The parameter *δ* represents a time scale for changes in the local actin polymerization speed.

The term Γ_*f,b*_ represents the friction term which changes with respect to the direction of motion of the cell’s edge, and is given by

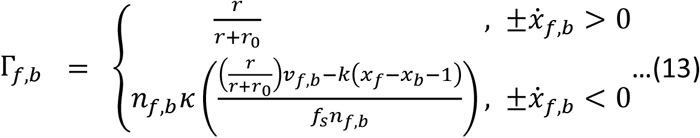

such that the top function in (13) applies to edges that extend outwards, while the complex lower function in (13) applies to edges that retract.

The terms 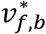 represent the steady state polymerization speed at the edges, and are coupled to the level of the polarity cue at the edge 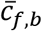 by

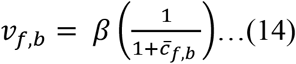

Where *β* is the maximal actin treadmilling flow, which couples the steady-state actin treadmilling flow to the saturated polarity cue at the cell’s edge. The exponential functional form of 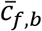 along the cell length is dominated by an advection-diffusion process along the cell, and is fully discussed in Maiuri, et al. 2015^[48]^.

For the construction of the phase diagram (Fig. 7B) we calculate two bifurcation curves: (1) The transition between the ‘no motility’ and the ‘smooth motion’ phases, calculated by finding a critical coupling strength *β*_*c*_ at which the actin polymerization speed is sufficient for the cell to become polarized. The value of *β*_*c*_ is determined by equating two polarization lengths which are derived from the model ^[4]^.

The first is a critical length of polarization due to the advection of the polarity cue

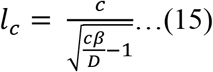

where *D* is the diffusion coefficient of the polarity cue, and *c* is a dimensionless quantity which encapsulates the concentration and its saturation properties ^[4]^.

The second critical length is derived from the force balance between the actin polymerization and the cell elasticity, and is given by

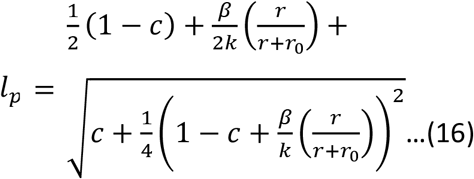

The analytical form of *β*_*c*_ as a function of *r* is given by

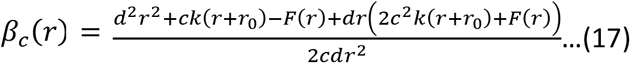

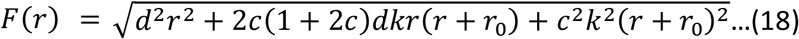

(2) The second transition line between the smooth motion and the stick slip motion is a Hopf bifurcation transition line which is obtained using a continuation method with AUTO07P ^[55]^. To calculate the persistence time (Fig. 7I) we first added noise to the system (Eqs.10-12). The noise is added to the actin polymerization speed in the equations, and its value was chosen to be Δ*v* = 2 (in dimensionless units, as *β*) to provide sufficient fluctuations for the cell to change its direction. The formula for the persistence time is given by

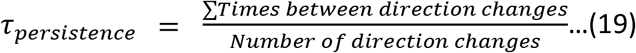

Throughout the simulations the fixed parameters that were used in the model are: *y* = 4, *D* = 4, *k* = 1, *f*_*d*_ = 5, *κ* = 20, *r*_0_ = 1 and *δ* = 100, *r* = 5. For Figure 7C-F the values that were used for the maximal actin retrograde flow are: *β* = 4,8,12,16 (with respect to (i-iv) in Figure 7B).

### 5.10 Actin flow measurements

The Halo-tagged actin was introduced into the cells by retrovirus and the cells were labeled by 5 nM Halo-TMR (Promega) for 1 hr ^[56]^. After replacing the medium and culturing for 4 hrs, the cells were imaged with the confocal microscope (FV1000, Olympus, with High-Sensitivity Detector). To determine the velocity of the retrograde flow, time-lapse image sequences were analyzed by kymographs using ImageJ. Moving actin features were visualized in the kymographs as streak lines. The velocity of the flow was then obtained from the slopes of these lines (shown by white solid lines in Supplementary Figure S10).

### 5.11 Statistics

Statistical analysis of the data was conducted using RStudio (RStudio, Boston, MA) software. The Shapiro-Wilks normality tested was used to check the normality of the data sets. Analysis of variance (ANOVA) was used to test for the statistical significance between different data sets. The following symbols are used to represent the statistical significance levels: *<0.05, **<0.01 and ***<0.001. If there is no comparison shown between any data sets, it implies that there is no statistically significant difference between them. All the error bars represent standard error of mean. Data was acquired from multiple, independent sets of experiments.

## Supporting Information

Supplemental Figures S1-S14. Movies 1-6.

Supporting Information is available from the Wiley Online Library or from the author.

**Supplementary Movie M1:** (Top) A WT cell spreading on a 135 nm diameter suspended fiber from an initial rounded state to a stretched state. (Bottom) An IRSp53 KO cell spreading on a 135 nm diameter suspended fiber from an initial rounded state to a stretched state. Time stamp is in hours:minutes.

**Supplementary Movie M2:** (Top) An IRSp53 WT cell spreading on a flat surface from an initial rounded state to a stretched state. An IRSp53 KO cell spreading on a flat surface from an initial rounded state to a stretched state. Time stamp is in hours:minutes.

**Supplementary Movie M3:** Protrusions of interest (shown by white arrows) along a 500 nm diameter suspended fiber (horizontal direction) for WT (top) and IRSp53 KO (bottom) cells. Note, the main cell-body is constrained along the large diameter (2000 nm) fiber in the vertical direction. Time stamp is in minutes.

**Supplementary Movie M4:** *Coiling* at the tip of protrusions (shown by white arrows) along suspended 135 nm diameter fibers (horizontal direction) for WT (top) and IRSp53 KO (bottom) cells. Note, the main cell-body is constrained along the large diameter (2000 nm) fiber in the vertical direction. Time stamp is in seconds.

**Supplementary Movie M5:** Cell force measurement using Nanonet force microscopy (NFM) for WT (top) and IRSp53 KO cell (bottom). In these videos, 500 nm diameter fibers (horizontal direction) are suspended on top of large diameter (2000 nm) fibers (vertical direction). Cells exert forces by tugging on fibers. Time stamp is in minutes:seconds.

**Supplementary Movie M6:** (Top) A WT cell migrating on a single 500 nm diameter suspended fiber. (Bottom) A KO cell migrating on a single 500 nm diameter suspended fiber. Centroids of cell body and nucleus are shown by red and green dots, respectively. Time stamp is in hours:minutes.

## Acknowledgment

We thank Prof. Masahito Ikawa (Osaka University) and Prof. Taro Kawai (Nara Institute of Science and Technology) for the CRISPR/Cas9 system, Prof Naoyuki Inagaki, and Dr. Takunori Minegishi for Halo-tagged actin, and all the members of the laboratories for technical assistance and helpful discussions. This work was supported by grants from the JSPS (KAKENHI JP20H03252) and JST CREST (JPMJCR1863) to S.S. N.S.G. is the incumbent of Lee and William Abramowitz Professorial Chair of Biophysics, and this research was supported by the Israel Science Foundation (Grant No. 1459/17). ASN acknowledges partial funding support from National Science Foundation (NSF, Grant No. 1762468). ASN and BB acknowledge the Institute of Critical Technologies and Science (ICTAS) and Macromolecules Innovative Institute (MII) at Virginia Tech for their support in conducting this study.

## Author Contributions

ASN conceived and supervised the research. ASN, SS, BB, and NG designed research. AM conducted experiments. HTH, TN, KHS, and SS generated the specific cell lines and performed actin flow measurements. JR and NG developed and implemented the theoretical model of 1D cell migration. AM, ASN, BB, SS, JR and NG analyzed data. AM wrote the manuscript. All authors contributed to editing of the manuscript.

Received: ((will be filled in by the editorial staff))

Revised: ((will be filled in by the editorial staff))

Published online: ((will be filled in by the editorial staff))

---

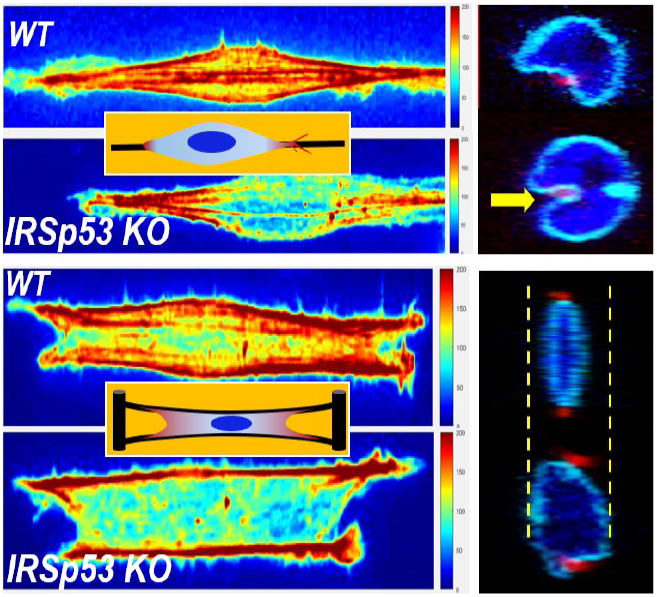

Using 1D suspended fibers capable of recapitulating 3D migration, we demonstrate that IRSp53 links protrusions to the nucleus through force generation to control cell migration in 3D. IRSp53 loss causes reduced actin dynamics at the tips of cells and a loss of actin stress-fiber-based contractility that transitions the stick-slip migration to a sliding-slower migratory mode due to uncoupling the nucleus from the cell body.

## Supporting Information

**Supplementary Figure S1:**
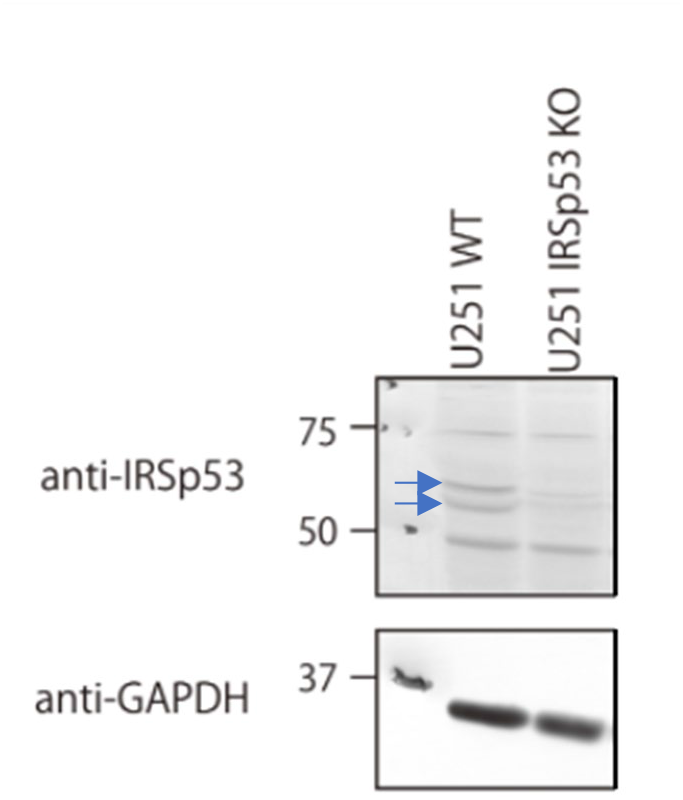
Western-blot demonstrates IRSp53 knock-out in U-251 cells. Bands of IRSp53 are shown by blue arrows.

**Supplementary Figure S2:**
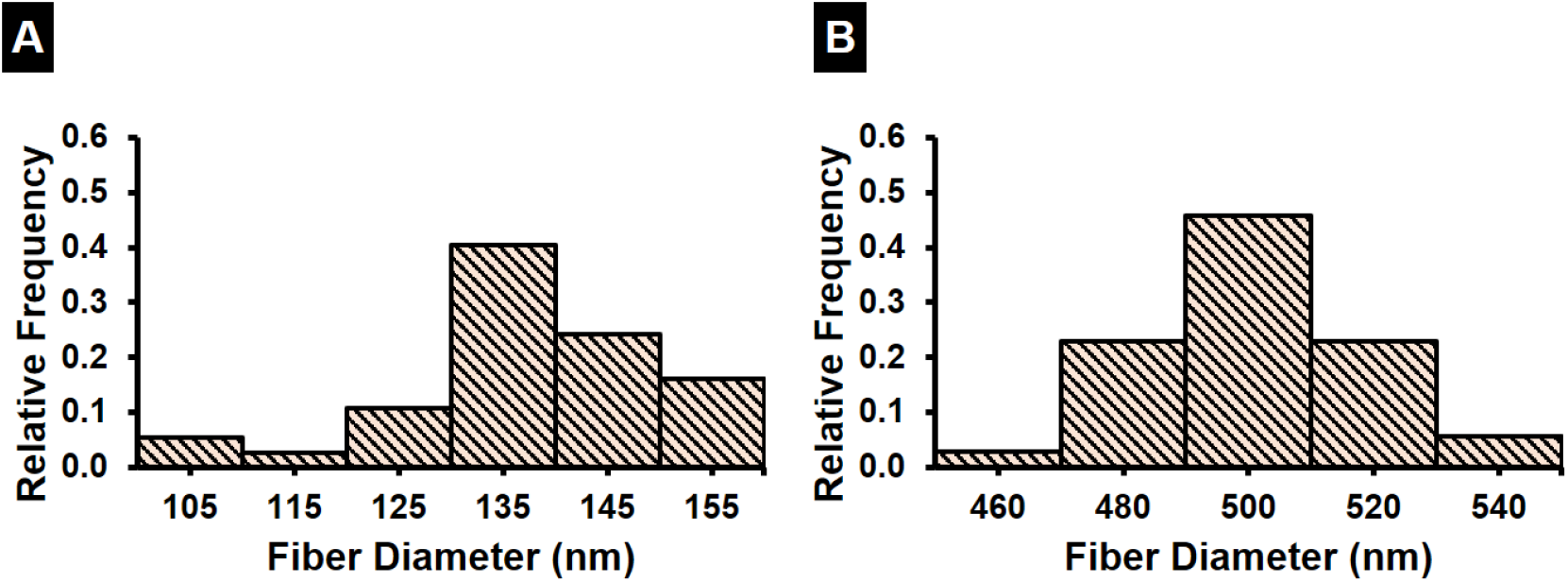
Quantifying fiber diameter distribution. Histograms showing the fiber diameter distribution for (A) ∼135 nm diameter, (B) ∼500 nm diameter fibers used in this study. n values are 40, and 35, for the ∼135 nm, and 500 nm diameters respectively.

**Supplementary Figure S3:**
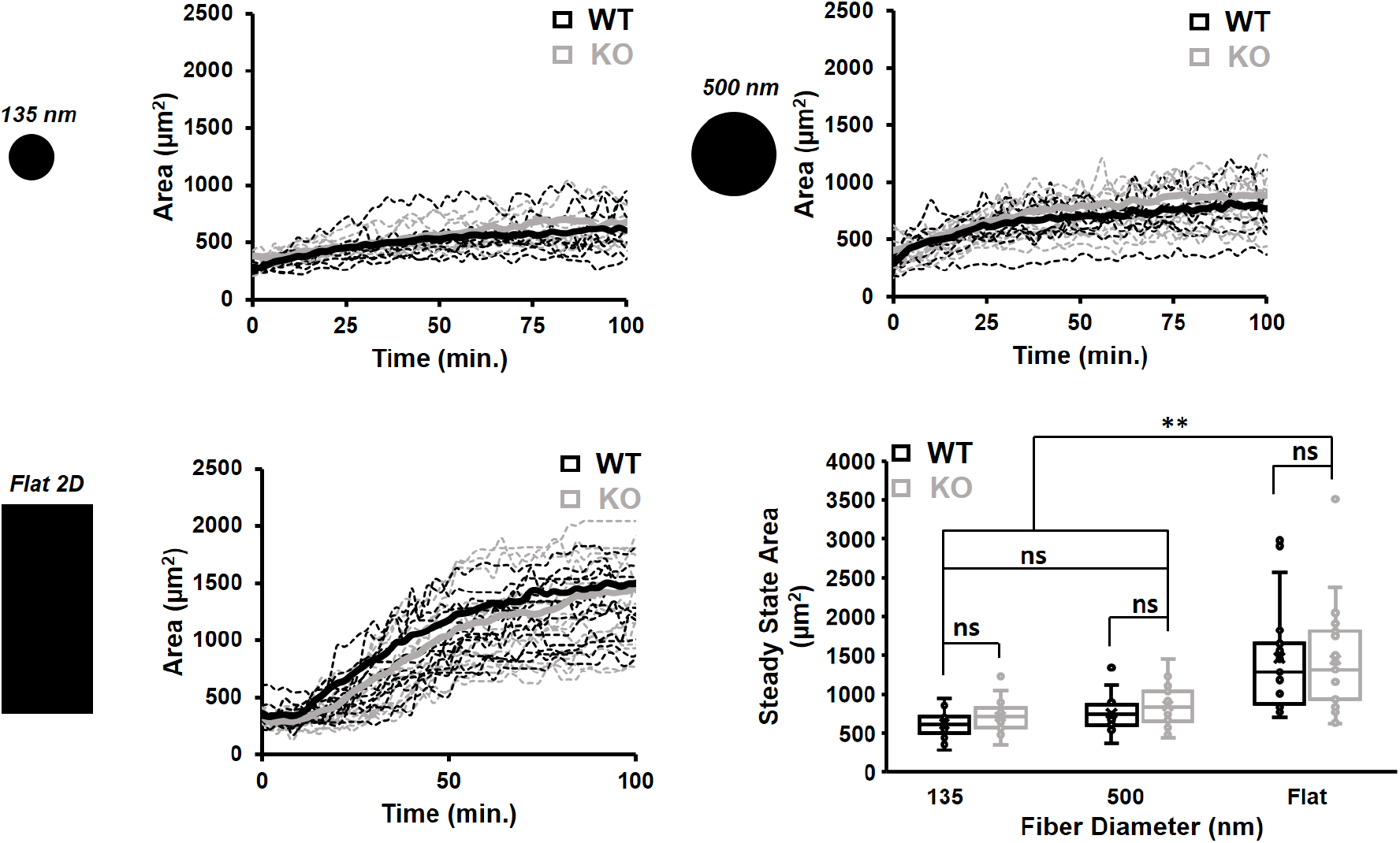
Cell Spreading on suspended fibers. Area calculations as cells (WT (black) and IRSp53 KO (grey)) spread on suspended fibers and flat control 2D. Steady state cell area on fibers is less than 2D. n = 20 for each cell category. All error bars shown represent standard error of mean.

**Supplementary Figure S4:**
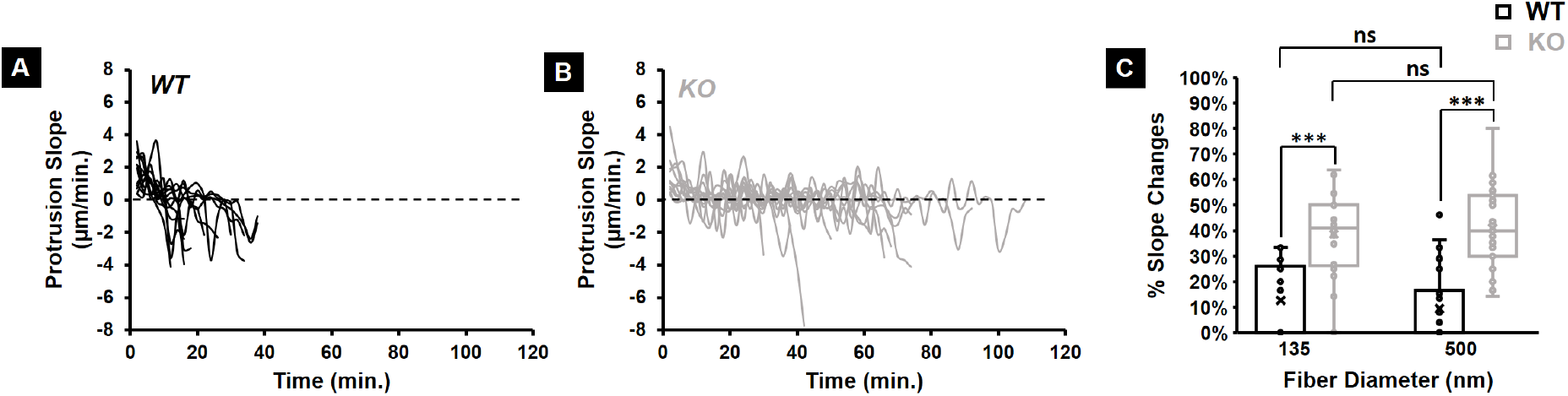
Quantifying the fluctuations in protrusion cycle for both KO and WT cells. Representative protrusive cycle slopes for both (i) WT and (ii) KO cells (12 profiles are shown here for each cell type on 135 nm diameter protrusive fibers) showing significantly more fluctuations in the KO case. (C) Quantification of the percentage of slope changes for WT and KO cells on both 135 nm and 500 nm diameter protrusive fibers. n values are 27 for both WT and KO cells on each fiber diameter. All error bars shown represent standard error of mean.

**Supplementary Figure S5:**
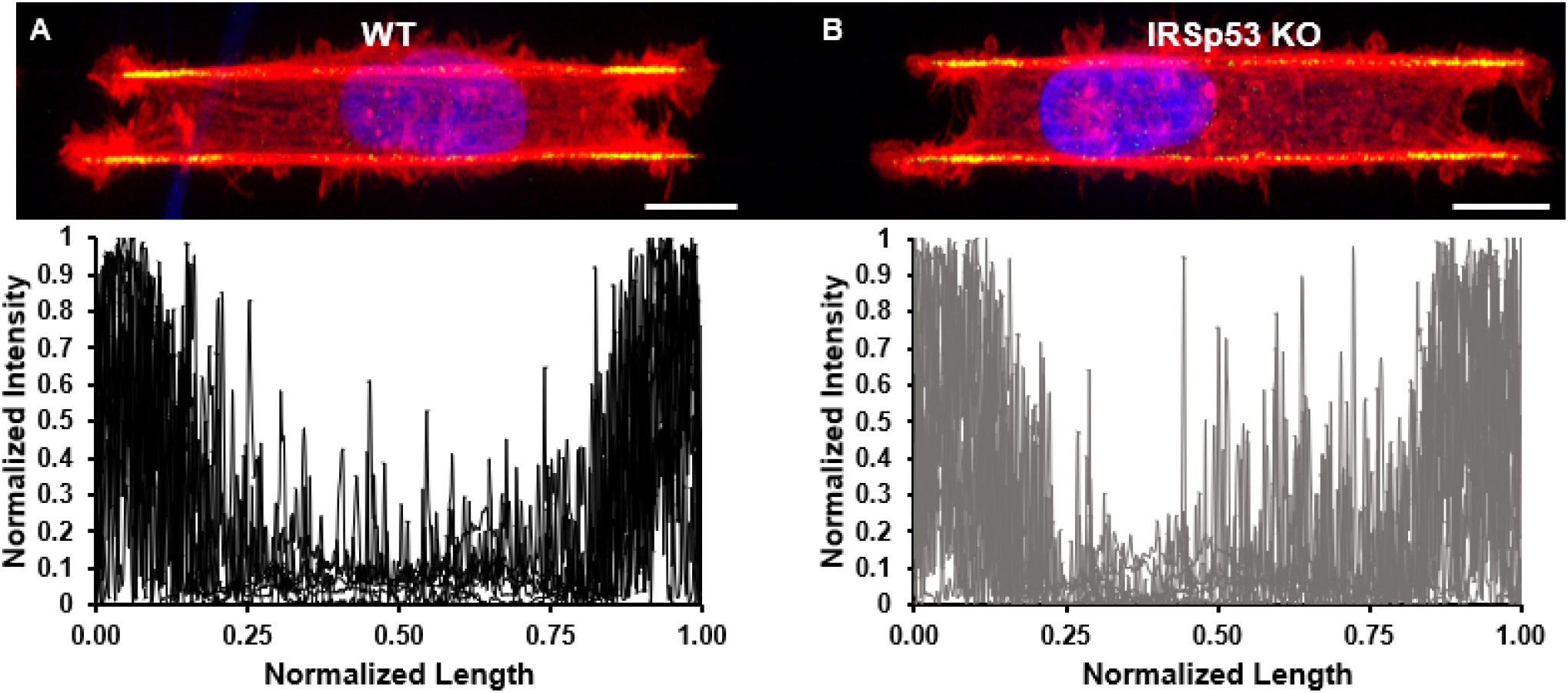
Quantifying the Paxillin distribution for both WT and KO cells on suspended parallel fibers. Normalized intensity profiles for Paxillin distribution along the cell length for (A) WT and (B) KO cells elongated on suspended parallel fibers. High intensity at the ends of the cell lengths indicates the clustering of Paxillin into focal adhesion clusters (FACs) at the cell edges (yellow). Profiles are obtained from 15 cells for each category. Scale bars are 10 μm.

**Supplementary Figure S6:**
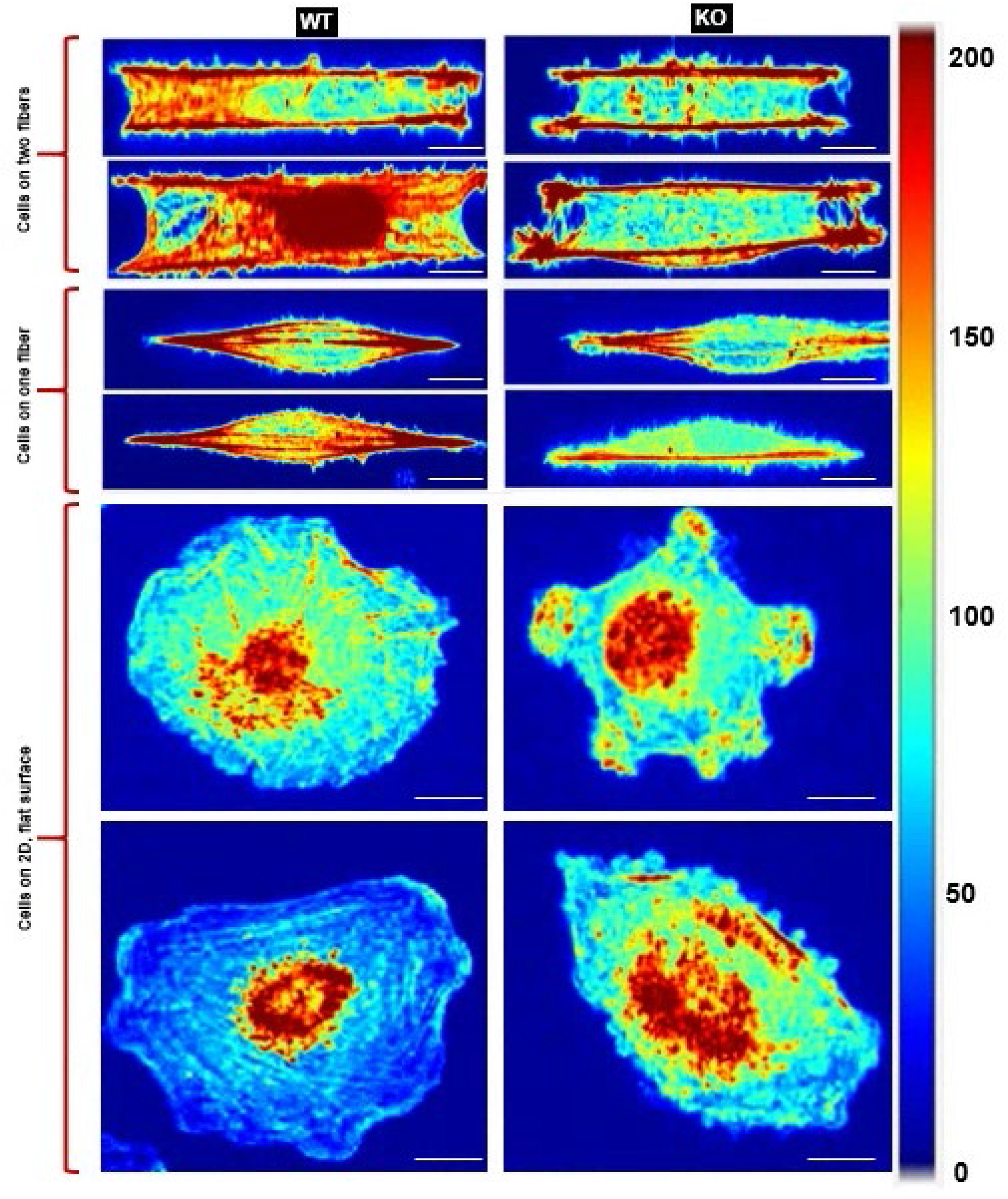
Actin stress fiber distribution. Color maps (arbitrary units) showing the actin stress fiber distribution for both WT and KO cells on 500 nm diameter suspended fibers in cells attached to two fibers (top two panels), single fibers (middle two panels) and on flat, 2D (bottom two panels). Scale bars are all 10 μm. In all cell shapes, KO cells have reduced number density of actin stress fibers.

**Supplementary Figure S7:**
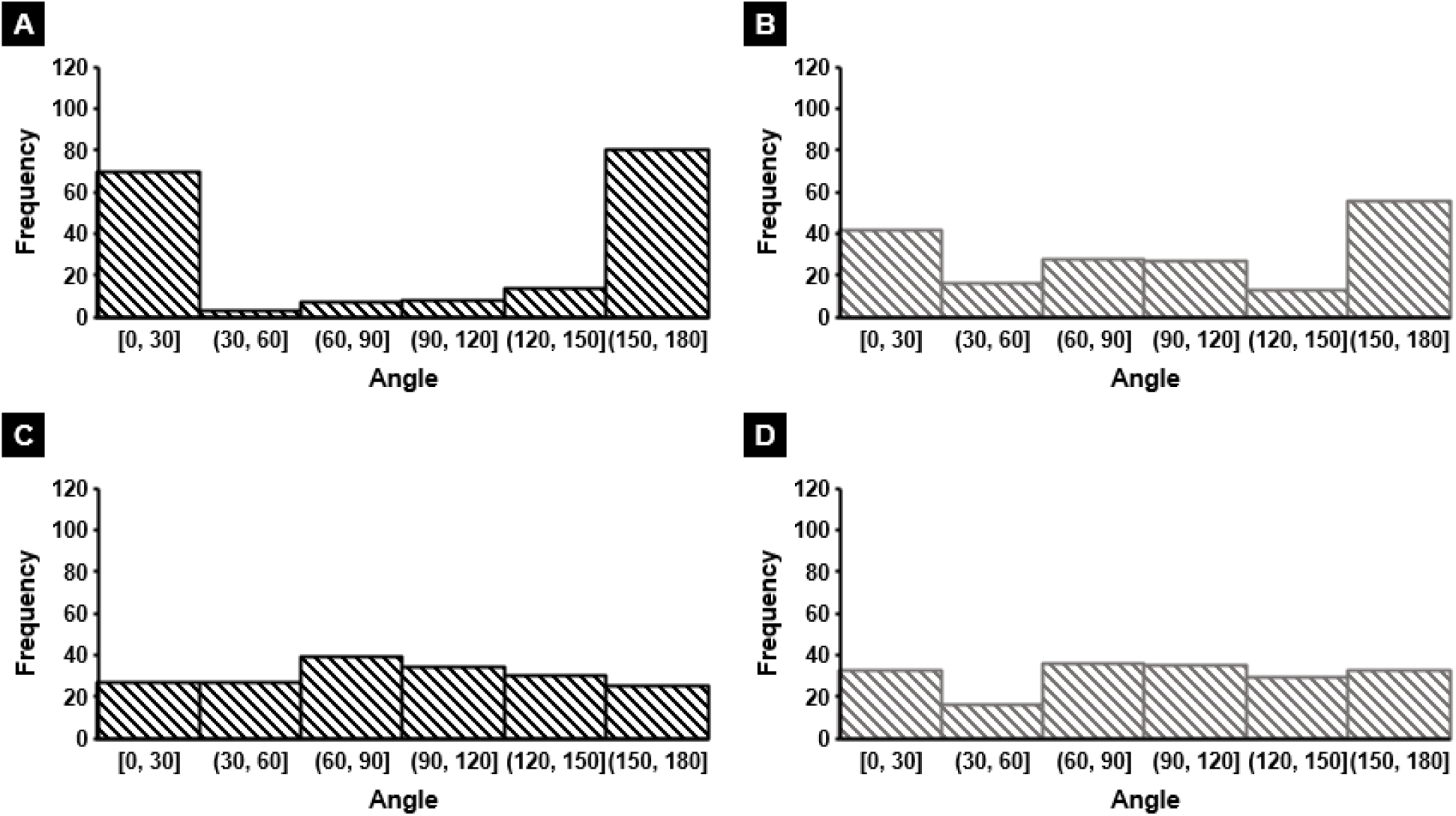
Stress fiber angular distribution. Histograms depicting the angular distribution of stress fibers for WT and **(B)** KO on parallel suspended fibers and **(C)** WT and **(D)** KO on flat 2D. n = 180 fibers for each category.

**Supplementary Figure S8:**
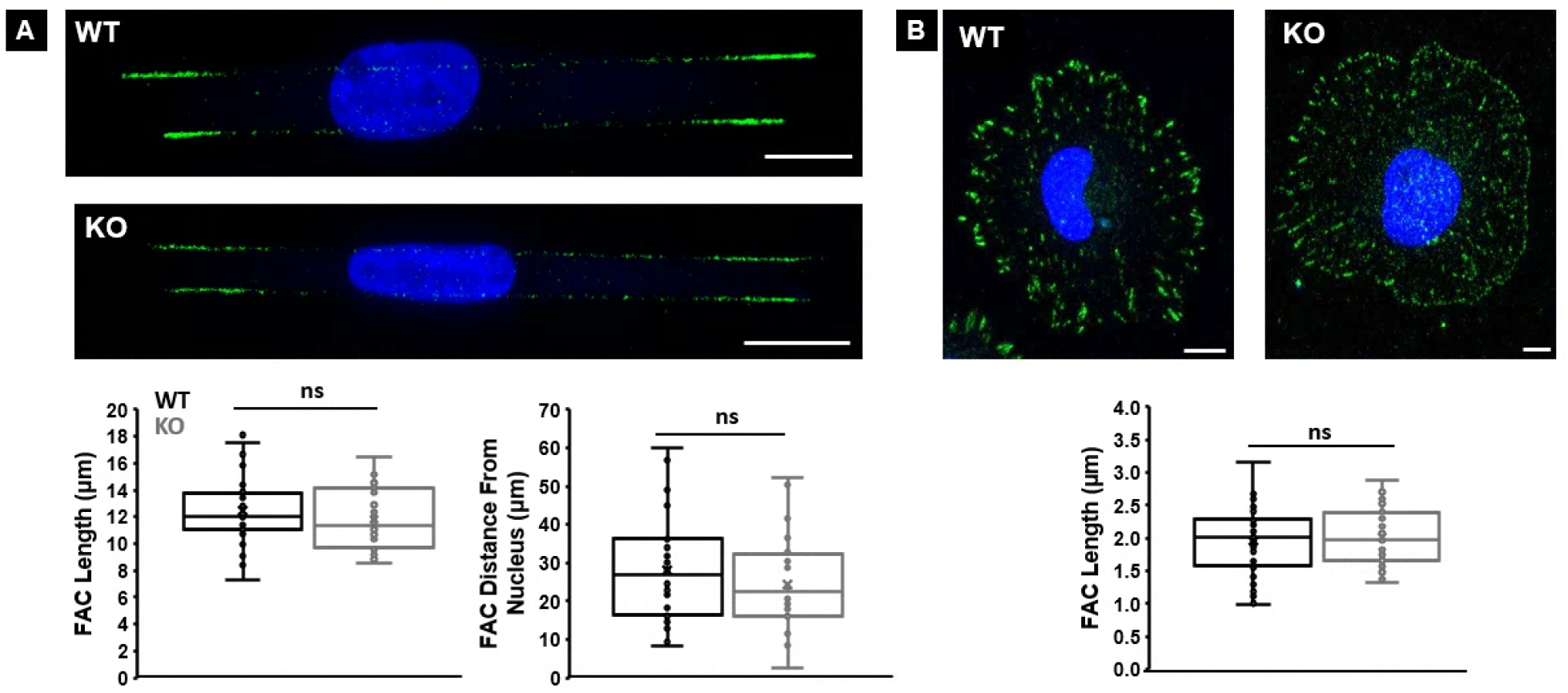
Focal Adhesion Cluster (FAC) distribution. Representative fluorescent images of paxillin clustering and associated FAC analysis on (A) 500 nm diameter parallel fibers and (B) flat, 2D for both cell types. Note that the FAC Distance From Nucleus metric is not applicable on flat, 2D surface (see Methods for more details). n values are 30 each for WT and KO cells on both fibers and on flat. All error bars shown represent standard error of mean.

**Supplementary Figure S9:**
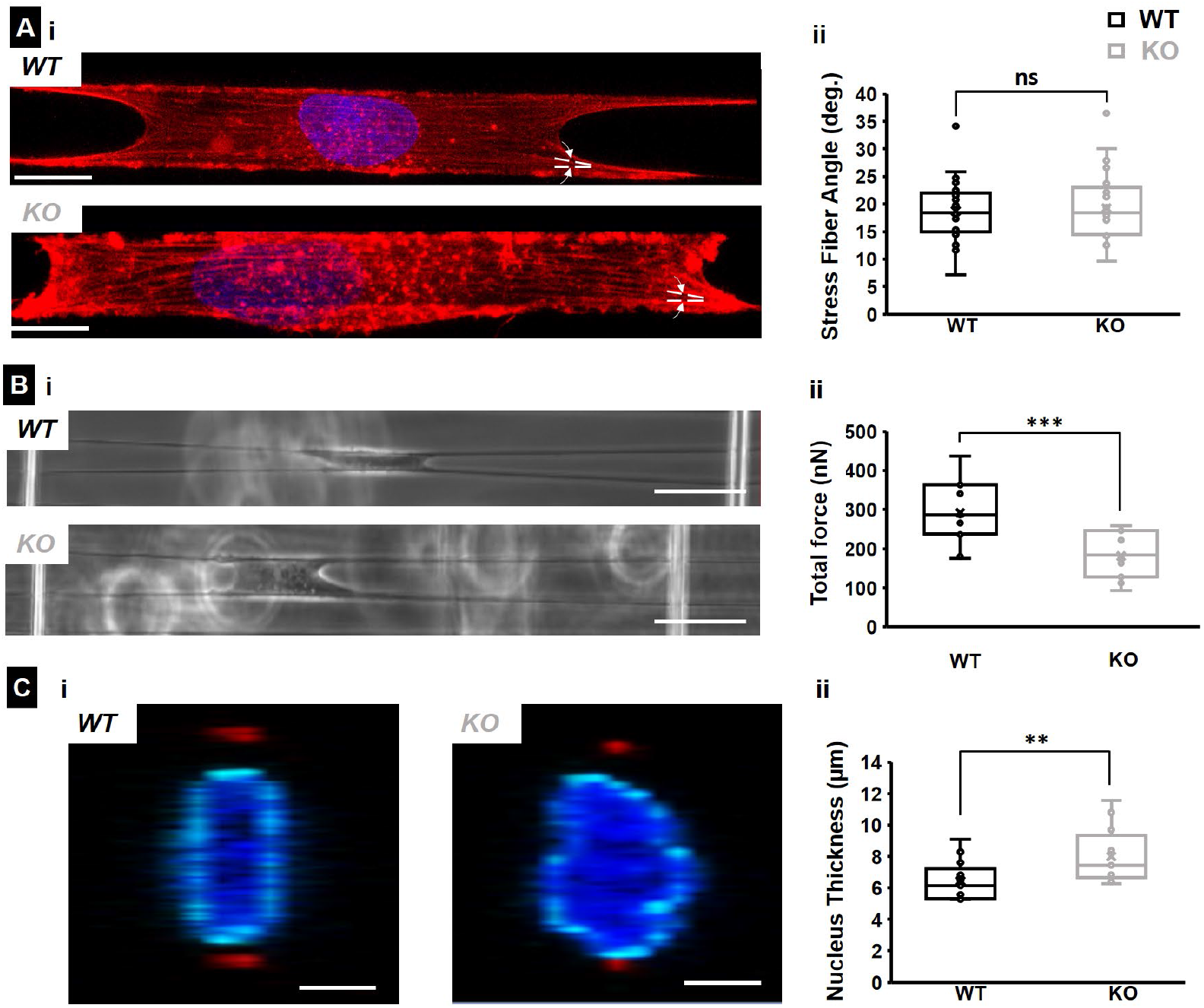
IRSp53 KO oral cancer cells show reduced forces and thicker nuclei compared to WT counterparts. (A) Representative fluorescence microscopy images of (i) WT (top) and KO (bottom) cells with f-actin stained in red and (ii) quantification of the stress fiber angles for both cell types on 500 nm diameter fibers. Scale bars are 10 μm. Dotted white lines in the fluorescent images depict the stress fiber angles. n values are 24 and 25 for the KO and WT cells respectively. (B) Representative phase images of (i) WT (top) and KO (bottom) cells exerting forces by pulling on suspended fibers and (ii) quantification of the forces exerted for both cell types on 500 nm diameter fibers. Scale bars are 50 μm. n values are 15 for both cell types. (C) Representative confocal images of (i) WT and KO nucleus cross-section (yz plane) on 500 nm diameter suspended fibers and (ii) quantification of the nucleus thickness. In the confocal images, the nucleus is in blue, the nuclear envelope is in cyan and the cross-section of the suspended fibers is in red. The yellow dotted lines depict the nucleus width. n values are 15 for both cell types. All error bars shown represent standard error of mean.

**Supplementary Figure S10:**
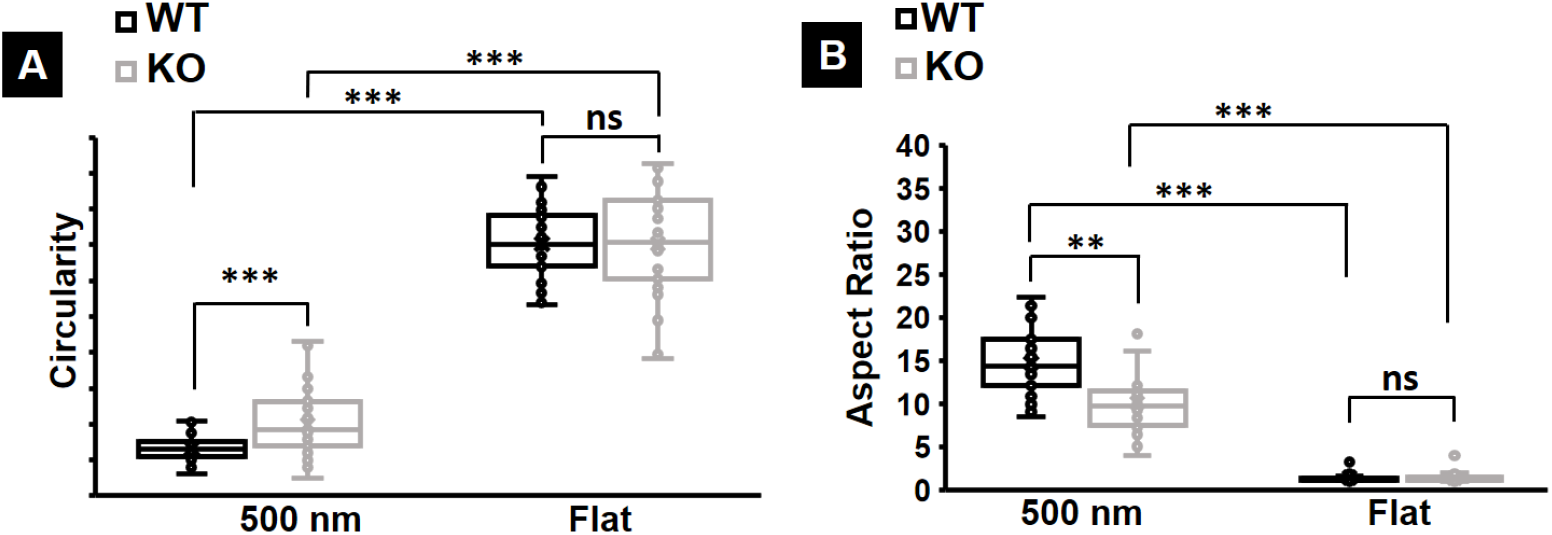
Quantification of average morphology metrics during migration. Average circularity and (B) aspect ratio of migrating WT and IRSp53 KO cells on 500 nm diameter suspended fibers and on flat, 2D surface. n for cells on fibers is 35 for both categories and on fibers is 30 for both categories. All error bars shown represent standard error of mean.

**Supplementary Figure S11:**
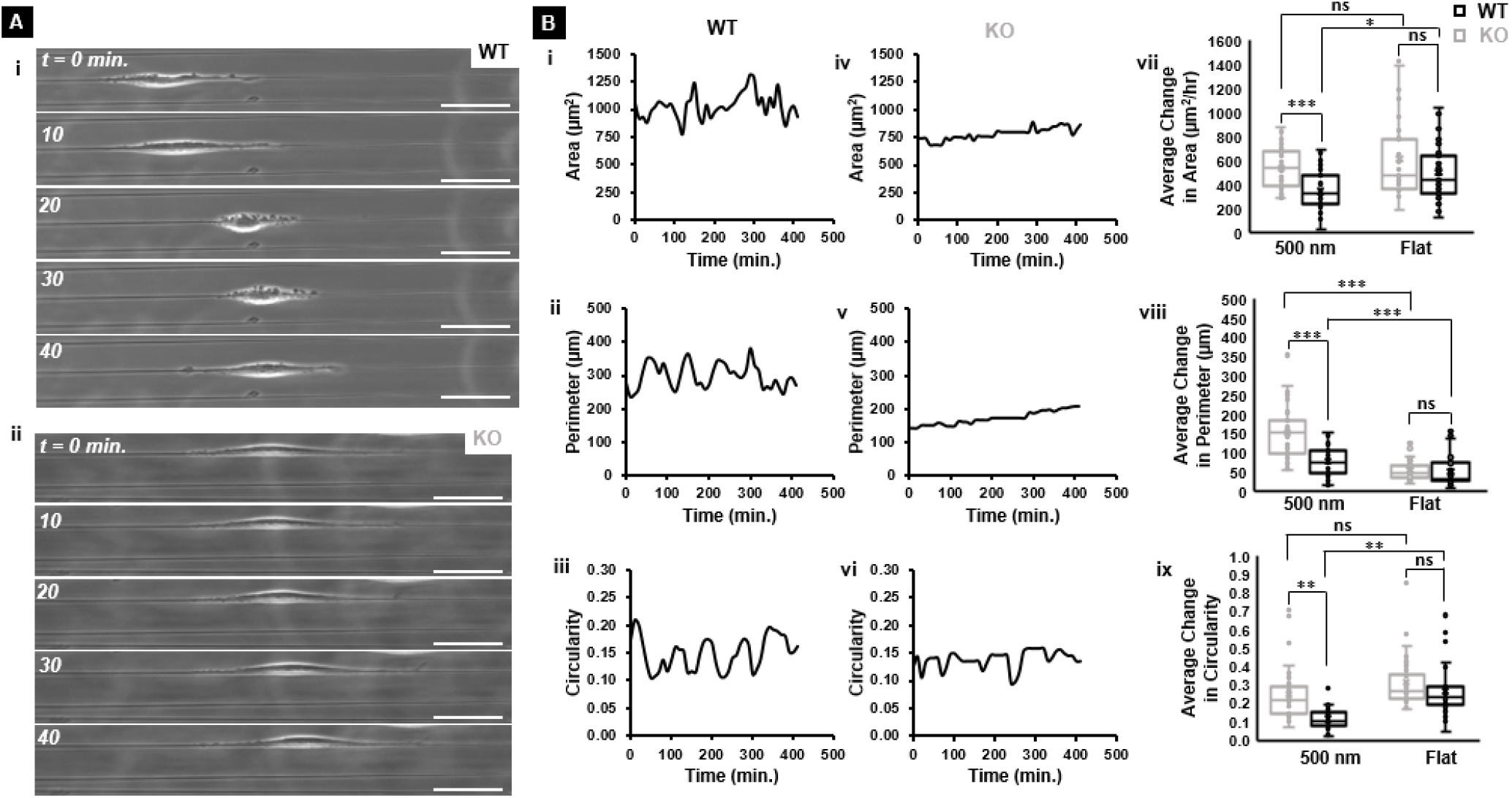
Quantification of the morphology changes during migration. (A) Representative phase images showing the change in cell shape during migration for (i) WT and (ii) KO cells. All scale bars are 50 μm. (B) Representative area, perimeter and circularity profiles for (i-iii) WT cells and (iv-vi) KO cells on 500 nm diameter suspended fibers. Quantification of the (vii) average change in area, (viii) average change in perimeter, and (ix) average change in the circularity of WT and KO cells on both 500 nm diameter suspended fibers and flat, 2D surfaces. n values are 35 for both cell types on the 500 nm diameter suspended fiber and 30 for both cell types on the flat surface respectively. All error bars shown represent standard error of mean.

**Supplementary Figure S12:**
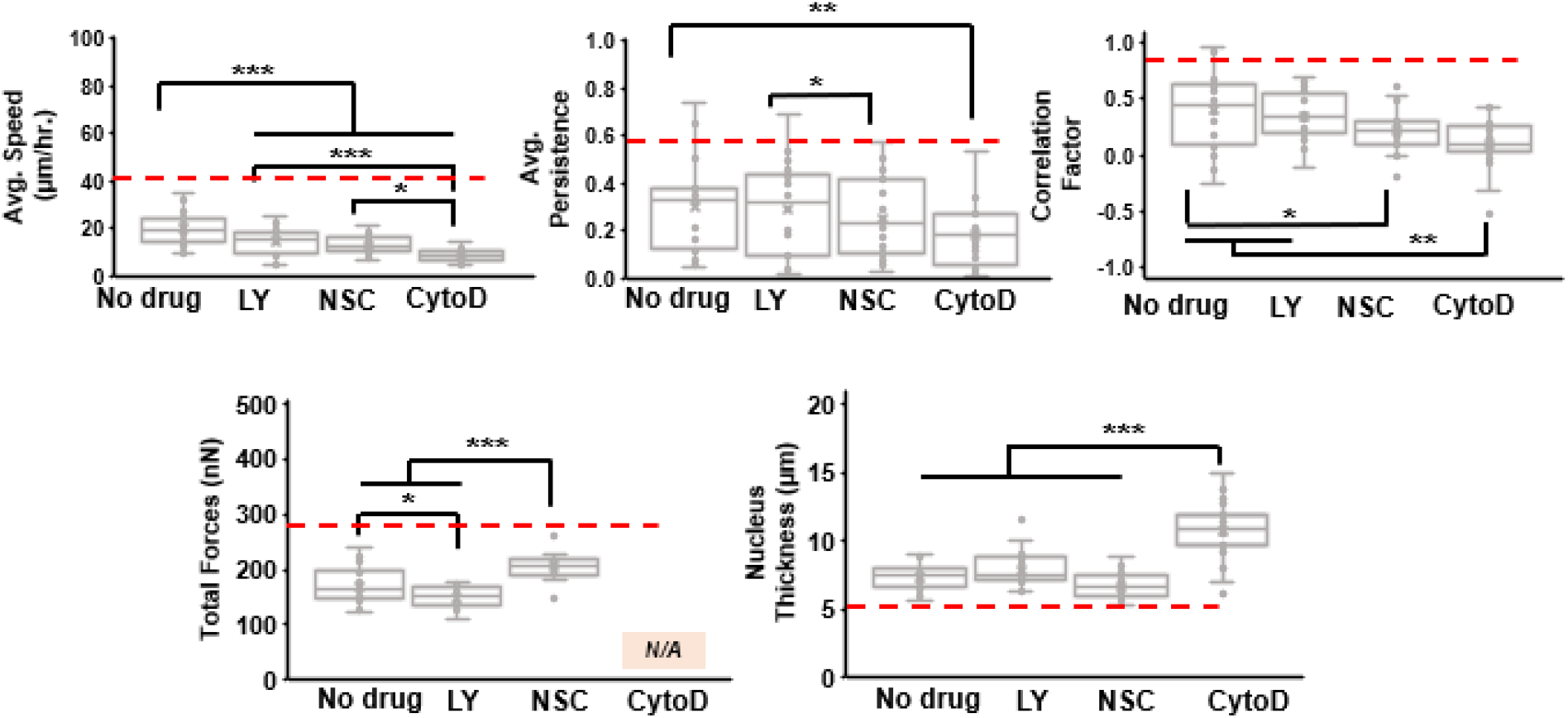
Pharmacological studies on IRSp53 KO cells. KO cells treated with LY294002 (PI-3 kinase), NSC23766 (Rac1), and Cytochalasin D (Actin). Red dashed lines represent the IRSp53 WT data without any drug treatment.

**Supplementary Figure S13:**
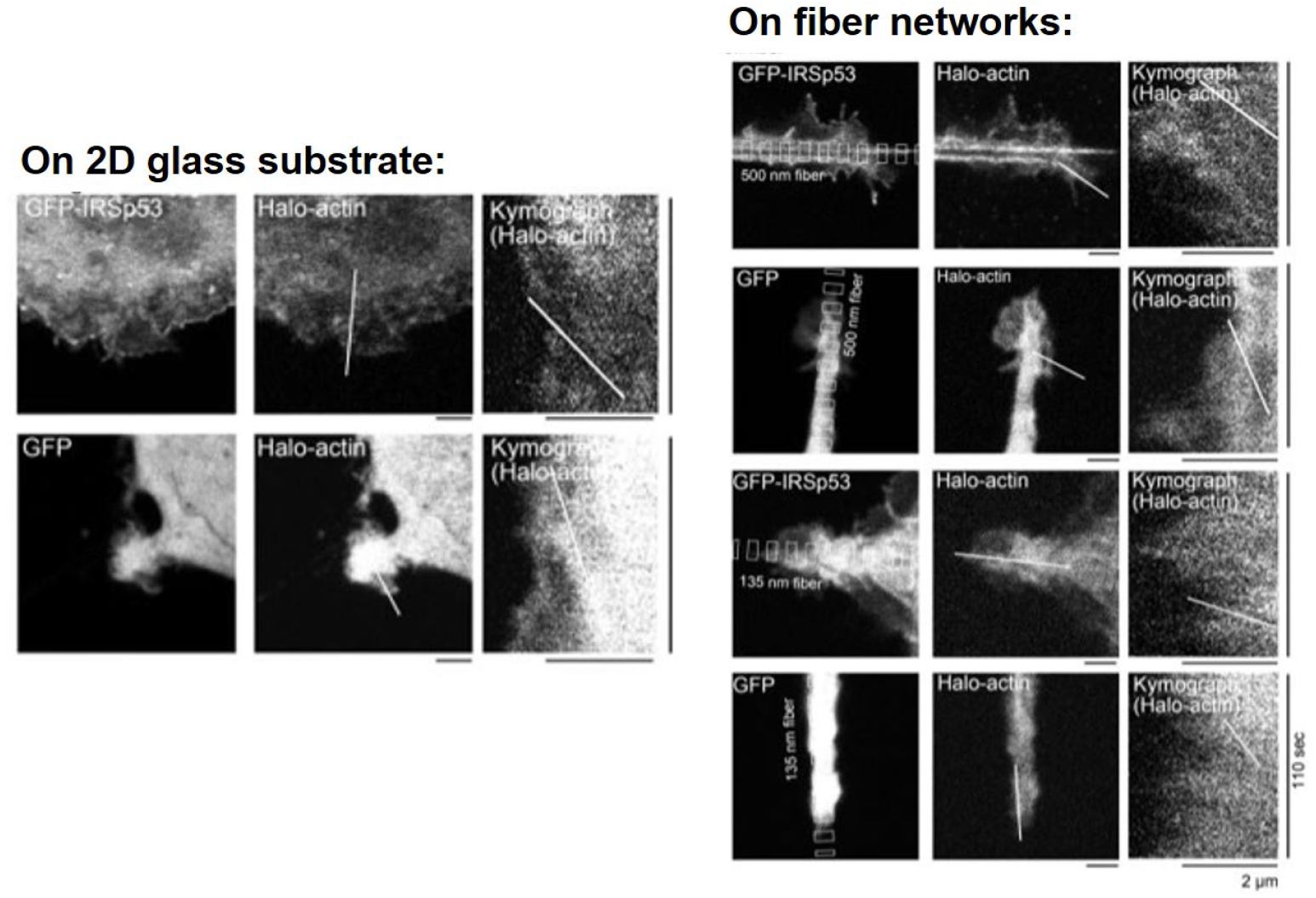
Speckle microscopy of actin flow. Fluorescence speckled microscopy images of both WT and IRSp53 KO cells on both 2D glass substrate (left panel) and suspended fiber networks (right panel).

**Supplementary Figure S14:**
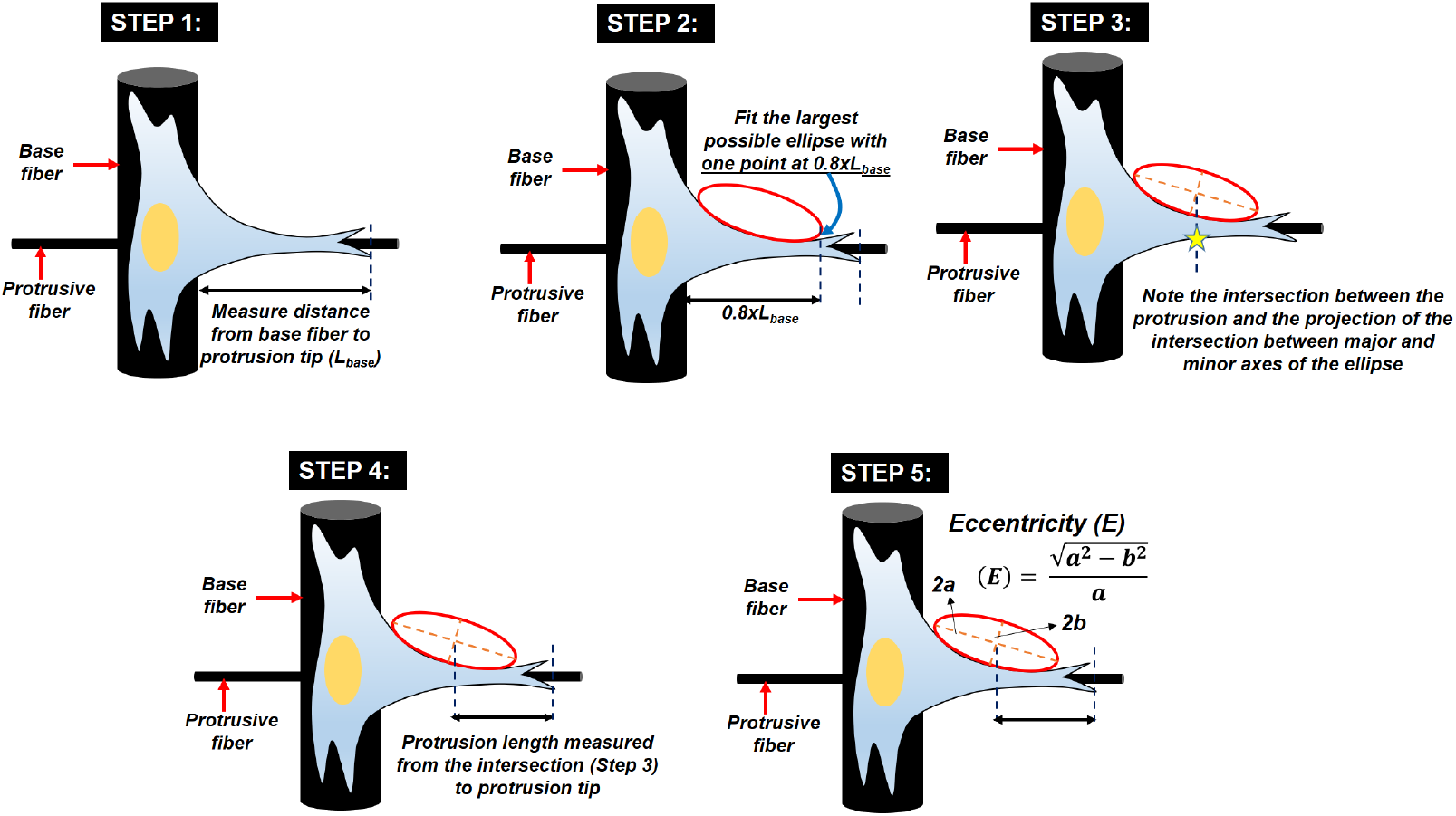
Protrusion measurements on suspended fiber networks. Schematic showing stepwise how the protrusion length and eccentricity are measured for single cells on our suspended fiber networks.

